# Right Inferior Frontal Gyrus Mediates Emotion–Cognition Interaction via Frequency-Specific Top-Down Modulation

**DOI:** 10.1101/2023.03.17.533197

**Authors:** Anya Dietrich, Edoardo Pinzuti, Yuranny Cabral-Calderin, Florian Müller-Dahlhaus, Michael Wibral, Oliver Tüscher

**Author notes:** These authors contributed equally to this work (Michael Wibral, Oliver Tüscher).

## Abstract

Interactions between emotional and cognitive processing play a critical role in determining human behavior. The ability to efficiently control these interactions is essential for goal-directed behavior. Previous research on emotion-cognition interaction in humans has primarily focused on its neural topography using techniques with either low temporal resolution (fMRI) or low spatial resolution (ERP), leaving the underlying neuronal mechanisms and their behavioral implications poorly understood. Here, we used EEG recordings from a large cohort of human participants (N=121) and employed source-reconstruction with finite-element head modeling during an emotional Flanker task. Our findings reveal a neural interaction between emotional interference and cognitive control within the pars triangularis of the right inferior frontal gyrus (rIFG). These results align spatially and conceptually with meta-analytic evidence from fMRI studies on interference inhibition processes, supporting the idea that the rIFG is crucial for interference inhibition, regardless of the interfering information type. Notably, the neural interaction between emotion and cognition in the rIFG was transient, peaking during the transition from emotional to cognitive processing, and was most pronounced in the beta-frequency band. Furthermore, conditional Granger-causality analysis of information flow indicated that the pars triangularis of the rIFG communicated with other functional subdivisions of the rIFG and parieto-occipital areas (such as the Precuneus and V2) in a frequency-specific manner. This information flow predicted individual emotional and cognitive interference effects on behavior. Overall, our findings highlight the functional and temporal segregation and integration of emotion, cognition, and their interaction in the rIFG. Furthermore, rIFG’s top-down modulation of visuo-attentional areas, ultimately shaping behavioral performance, underscores the importance of stimulus interference inhibition for emotion-cognition interaction.

**Author summary:** Understanding the interplay between emotion and cognition is crucial for understanding human behavior. Our study investigates the role of transient oscillatory activity using source-localized EEG in a large cohort. We identify a spatial overlap of the interaction of emotion and cognition processing in the rIFG’s pars triangularis, aligning closely with localizations indicated by meta-analytic fMRI evidence.

This interaction, primarily in the beta-frequency (around 20Hz) band, is transient, peaking during the shift from emotional to cognitive processing. Granger-causality analysis reveals frequency-specific communication patterns within the rIFG and parieto-occipital areas, predicting individual emotional and cognitive interference effects.

Our findings elucidate the functional and temporal dynamics of emotion-cognition interaction within the rIFG, providing insights into the nuanced neural processes shaping human behavior.

## Introduction

The interplay of emotional and cognitive processing is of fundamental practical importance to human behavior and psychopathology: emotions can drive us to excel and exceed ourselves, but also interfere disastrously with task performance to a degree that can change the course of our lives [1]. Due to the latter, emotional interference inhibition, i.e., the inhibition of distracting, salient emotional stimuli, is critical for well-being and mental health, especially as even non-task-relevant emotional stimuli receive preferred processing resources – at least when sufficiently arousing and salient [2–4]. Such emotional interference inhibition requires at least the integration of emotional with cognitive processing in the brain, and possibly the recruitment of additional neural systems that exert influence over both emotional and cognitive processing.

Emotional interference is thought to arise from a competition for neural resources between emotion processing and core cognitive functions, particularly updating, shifting, and inhibition [3, 5, 6]. However, the precise neural mechanisms governing this competition remain unclear. Specifically, it is not well understood whether the resolution of this competition is handled locally within the engaged neural networks or whether distinct brain regions exert additional control to suppress emotional interference. Addressing this question is crucial, as alterations in emotional and non-emotional interference processing are observed in various mental disorders, including major depression [7], bipolar disorder [8], and post-traumatic stress disorder [9]. Resolving these questions requires an approach that provides both high spatial and temporal resolution to map the neural circuits where emotion and cognition interact. To date, most studies have relied on functional magnetic resonance imaging (fMRI) for spatial precision or magneto/electroencephalography (M/EEG) for temporal precision, but not both. However, given that fMRI is limited in capturing real-time neural signaling, the only feasible solution at present is to maximize the spatial resolution obtained from M/EEG source reconstruction.

To this end, we performed a large-cohort (N=121) EEG study using an emotional Flanker task, combined with high-resolution anatomical MRI (T1 and T2 contrasts), tissue segmentation, Finite Element Modeling (FEM) of the electromagnetic forward solution, and beamforming for the inverse solution [10]. This multimodal approach allowed us to examine the emotion-cognition interaction at the neural level across time, space, frequency, and information transfer, thus achieving high temporal and spatial resolution. Based on prior evidence suggesting the role of beta-band oscillations in inhibitory control [11, 12] and the importance of top-down regulation in emotion-cognition interactions, we formulated the following hypotheses:

- We hypothesized that emotion-cognition interactions would be reflected in beta-band activity, indicating the role of inhibitory control over emotional interference.
- We hypothesized that inhibitory control would be exerted by frontal regions through local information flow and top-down modulation to regions associated with the behavioral interference effects.

Our study employed a whole-brain analysis approach to investigate emotion–cognition interactions, examining neural activity across all cortical sources rather than restricting analyses to predefined regions of interest. Interestingly, our findings revealed that the inferior frontal gyrus (IFG), particularly the right pars triangularis, was the only source exhibiting a significant interaction effect in the beta-band. Furthermore, Granger causality analysis demonstrated that the IFG exerts top-down modulation over posterior regions (e.g., Precuneus, V2), in a manner that directly impacts behavioral interference effects. This finding indicates that the IFG does not only resolve emotion-cognition interaction, but actively regulates neural activity in posterior regions to modulate interferences. These findings provide novel insights into how prefrontal dynamics resolve emotion-cognition competition and highlight the role of frequency-specific network communication associated with inhibitory control of interferences.

## Results

### Emotional interference versus emotional interference inhibition and their relation to statistical contrasts

The aim of this study was to elucidate the neuronal mechanisms of the critical interplay of emotional and cognitive processing. Here, the emotional Flanker task used specifically operationalizes this interplay by the (implicit) processing of visual emotional stimulation / information preceding a (non-emotional) cognitive control task (cognitive stimulus and response interference [13]). This emotional processing interferes with cognitive processing (as shown by a behavioral effect). However, this emotional interference needs to be *inhibited* to enable and maintain cognitive processing despite this emotional interference. As the combination of two concepts, i.e., emotional interference and emotional interference *inhibition*, together with an analysis of both behavioral and neural data can lead to some preventable misunderstandings, we start by explaining and defining two important aspects related to the interpretation of our results.

First, we speak of *emotional interference* at the behavioral level, when emotion processing interferes with the performance in the cognitive task, i.e., when there is a statistical *main effect of emotional valence* on task performance. In contrast, we speak of *emotional interference inhibition* when there is a statistical *interaction effect* between cognitive load and emotional valence. Strictly speaking, it would be better to talk about emotional interference “modulation” here, as inhibition indicates a reduction in emotional interference, whereas the interaction effect could be such that we’d see an amplification of emotional interference. We stick, however, to the term previously used in the literature and discuss the sign of the observed interaction instead – where necessary.

Second, at the level of neural activity, we have to be additionally aware of the fact that the statistical *main effect of emotional valence* may reflect emotional interference, as above, but also pure emotional processing. –

The aim of this study was to elucidate the neuronal mechanisms underlying the interplay between emotional and cognitive processing. We employed an emotional Flanker task designed to operationalize this interplay, in which (implicit) processing of visually presented emotional stimuli (emotional images) precede a cognitively demanding non-emotional Flanker task involving stimulus and response interference [13]. This task design enables us to investigate how emotional processing interacts with and potentially disrupts cognitive performance. The observed behavioral effects confirm that emotional processing interferes with cognitive task performance. However, to maintain cognitive function despite this interference, emotional interference must be *inhibited*.

Given the conceptual overlap between emotional interference, emotional interference *inhibition*, and their neural and behavioral correlates, it is important to establish clear definitions to ensure precise interpretation of our results. Below, we define two critical aspects of these constructs in relation to our statistical contrasts.

First, we define *emotional interference* at the behavioral level as the phenomenon where emotion processing disrupts cognitive task performance. This is statistically reflected in a *main effect of emotional valence* on task performance, indicating that certain emotional stimuli impair reaction times or accuracy independent of cognitive task demands.

In contrast, we define *emotional interference inhibition* as the process by which this interference is reduced or modulated. This is statistically represented by a significant *interaction effect* between cognitive load and emotional valence. While the term “inhibition” suggests a reduction in interference, an interaction effect does not necessarily indicate suppression; it may also reflect an amplification of interference under specific conditions. To maintain consistency with previous literature, we retain the term “emotional interference inhibition” but explicitly discuss the direction of effects where necessary.

Second, at the level of neural activity, we must differentiate between the *main effect of emotional valence* and its potential interpretations. While this effect may indicate emotional interference (as observed in behavior), it may also reflect pure emotional processing independent of cognitive demands. Thus, careful interpretation is necessary when linking neural activity to behavioral outcomes.

### Large-cohort, FEM-based EEG-beamforming for a highly resolved study of neural sources in emotion-cognition interaction

EEG data were collected from a large cohort of healthy human participants (n=121; age 18 to 54 years) while they performed an emotional Flanker task. During this task, emotional (negative/neutral) images preceded the Flanker stimulus (incongruent/congruent). EEG data were analyzed using a combination of finite element head modeling (FEM) for the electromagnetic forward solution, and beamformer EEG source localization, yielding temporally and spatially highly resolved source data (see Fig 1 A and the Task Design section for a detailed description. See also Fig 9 for the full analysis pipeline). We focused our analyses on the time window after onset of the Flanker stimulus, i.e., tROI Flanker in Fig 1, because only then an emotion-cognition interaction could occur. Results from these analyses are presented in the following sections. For an analysis of neural activity before Flanker stimulus onset, i.e., during the initial emotional processing following the display of the emotional images only (tROI Emo), please see supplementary material section Supplementary results of source localization and.

**Fig 1.**
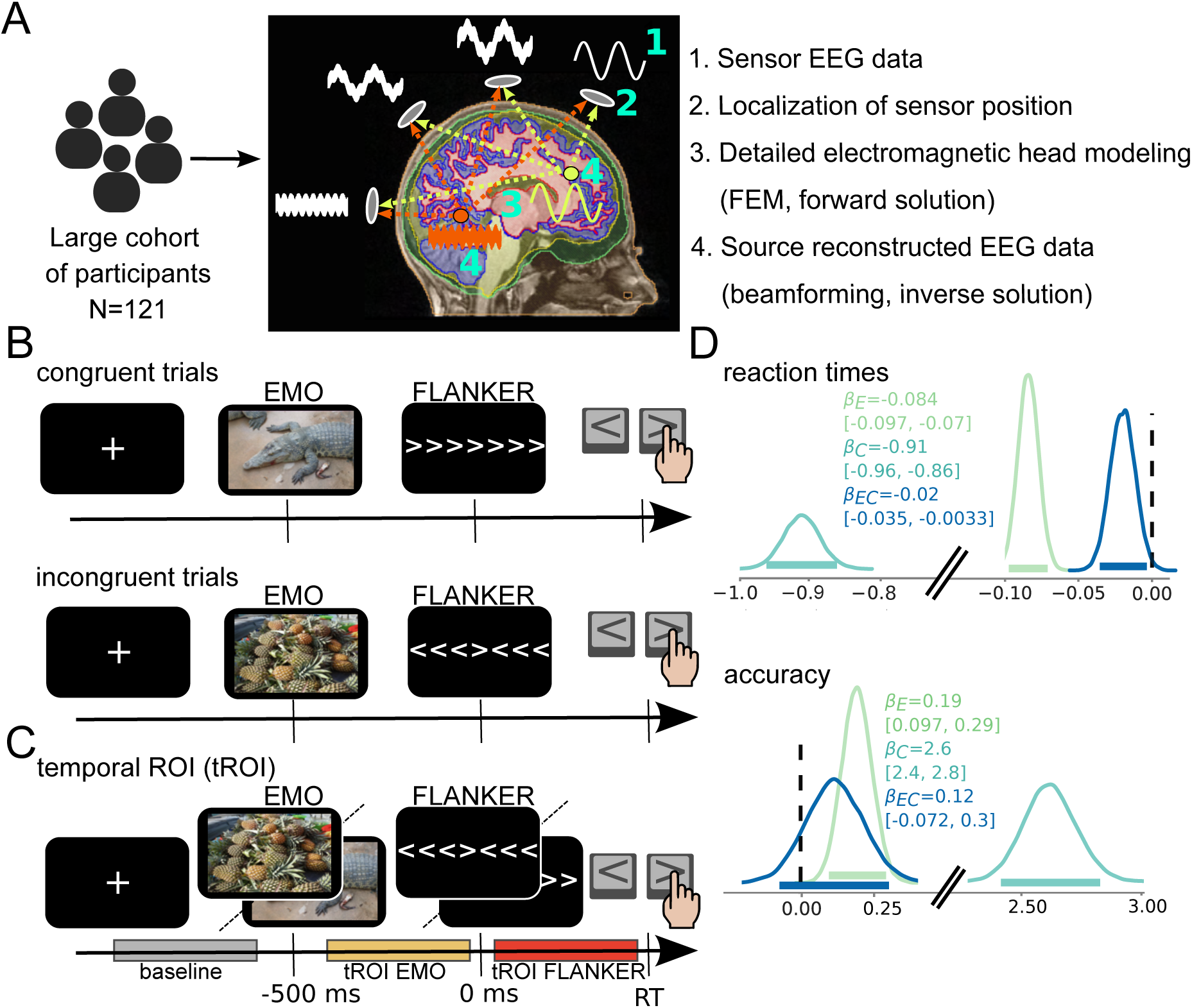
Emotional Flanker task design and behavioral effects. A, We collected EEG data (1) from 121 healthy human participants and located each electrode’s position (2). Using individualized Finite Element Models (FEM) of the head from structural MR images (3), we calculated electrical signal propagation in the brain (forward solution). With a beamforming algorithm, we reconstructed the signals in source space in the brain (4) – based on scalp recordings, sensor positions, and the forward solution (Please see Materials and Methods and Fig 9). B, Basic trial design: images with neutral or negative valence (placeholder images indicate IAPS pictures) precede onset of a congruent or incongruent Flanker stimulus (time, 0 ms), determining which button to press in the subsequent behavioral response (left for a central *<* symbol or right for a central *>* symbol). C, Temporal regions of interest (tROI) of 380 ms length tROI EMO (−400 to −20 ms relative to Flanker stimulus onset, yellow) during presentation of the IAPS picture, tROI FLANKER (50 ms to 430 ms, red) after Flanker cue onset and baseline (−1000 to −620 ms relative to Flanker cue onset, gray) were used for further source reconstruction. Of note, tROIs were defined for each frequency band separately, and a duration of 380 ms is an average value; see below, Task Design and Temporal ROI definition D, Posterior distribution of the Bayesian linear regression model parameters for reaction time (cognitive stimulus interference) and accuracy (cognitive response selection interference). *β*_meanpower_= *β* mean power, *β*_C_=cognition effect, *β*_E_=emotion effect and *β*_EC_=interaction effect. Inset values indicate the median and 94% credible intervals of the posteriors; the latter values are also represented by horizontal bars of the same color. Priors in.

### Behavioral effects demonstrate an emotion-cognition interaction

To quantify the behavioral interference effect of emotion on goal-directed cognitive processing, we used Bayesian Hierarchical linear models. In Bayesian regression models, the more plausible it is that experimental effects exist, the more exclusively positive or negative the posterior distribution of the regression parameters is [14]. For brevity of the presentation we therefore speak of an ‘effect’ here when the 94% posterior highest density interval (HDI) contains exclusively positive or negative values, but urge the reader to inspect the full posterior distributions to assess the available evidence and uncertainty. In this sense, reaction time (as a measure of cognitive stimulus interference) and accuracy (as a measure of cognitive response interference) revealed effects of emotion and cognition, as well as a statistical emotion-cognition interaction effect, as all posterior HDIs of the regression parameters were negative (reaction time) or positive (accuracy). As for the emotional effect, reaction times were slower (posterior median: *β_E_* = −0.084, posterior HDI: [−0.097, −0.07]) and accuracies lower (posterior median: *β_E_* = 0.19, posterior HDI: [0.097, 0.29]) for negative compared to neutral trials. As for the effect of cognition, reaction times were slower (posterior median: *β_C_* = −0.91, posterior HDI: [−0.96, −0.86]) and accuracies lower (posterior median: *β_C_* = 2.6, posterior HDI: [2.4, 2.8]) for incongruent compared to congruent trials, corroborating well-known cognitive interference effects in the Flanker task. Overall, the behavioral effect of cognition was larger than that of emotion and the statistical emotion-cognition interaction (see Fig 1, for detailed behavioral description see Supplementary information about behavioral effects of emotion-cognition interaction). A statistical interaction effect of emotion and cognition was observed for reaction time (posterior median: *β_EC_* = −0.02, posterior HDI: [−0.035, −0.0033]), with the effect of emotion (i.e., negative vs. neutral condition) being stronger during low compared to high cognitive load (i.e., congruent vs. incongruent condition). A possible interpretation is that a stronger cognitive load might use more processing resources than a lower cognitive load, thereby draw additional resources from emotional processing and as consequence reduce the emotional interference effect, i.e., the resource competition is internally resolved in favor of the more demanding process [3, 4].

For comparison with previous studies using frequentist methods, we also performed additional linear mixed effect analysis in the Supplementary information about behavioral effects of emotion-cognition interaction and, and obtained highly similar results (see Supplementary information about behavioral effects of emotion-cognition interaction and for comparison).

### Source-level activity reveals rIFG as a key region for emotion-cognition interaction

For the analysis of neural source activity during emotion-cognition interaction, i.e., during cognitive processing after emotional image presentation (tROI Flanker), we contrasted neural source-level spectral power in a non-parametric 2×2 repeated-measures cluster permutation ANOVA with the within-subject factors emotion (negative/neutral) and cognition (incongruent/congruent). The only source showing significant effects in all three contrasts was the rIFG (n=103, main effects of emotion (*pars orbitalis*, MNI peak coordinates x=45, y=40, z=0, F-Value=10.4, p*<*1.9996e-04), cognition (*pars opercularis*, MNI peak coordinates x=55, y=10, z=10, F-Value= 7.9, p*<*1.9996e-04), and interaction effect (*pars triangularis*, MNI peak coordinates x=45, y=40, z=20, F-Value= 9.6, p*<*1.9996e-04)) – with these effects occurring in the *β*-band (Fig 2). We additionally report results from exploratory analyses in all other frequency bands during tROI Emo and tROI Flanker in the supplementary material under Supplementary results of source localization. An overview of all sources, with their respective MNI peak coordinates, can be found in Table 1 for the *β*-band and in for the other frequency bands; also see (,) for *θ*-, () for *γ*- () and for high *γ*-band surface plots.

**Fig 2.**
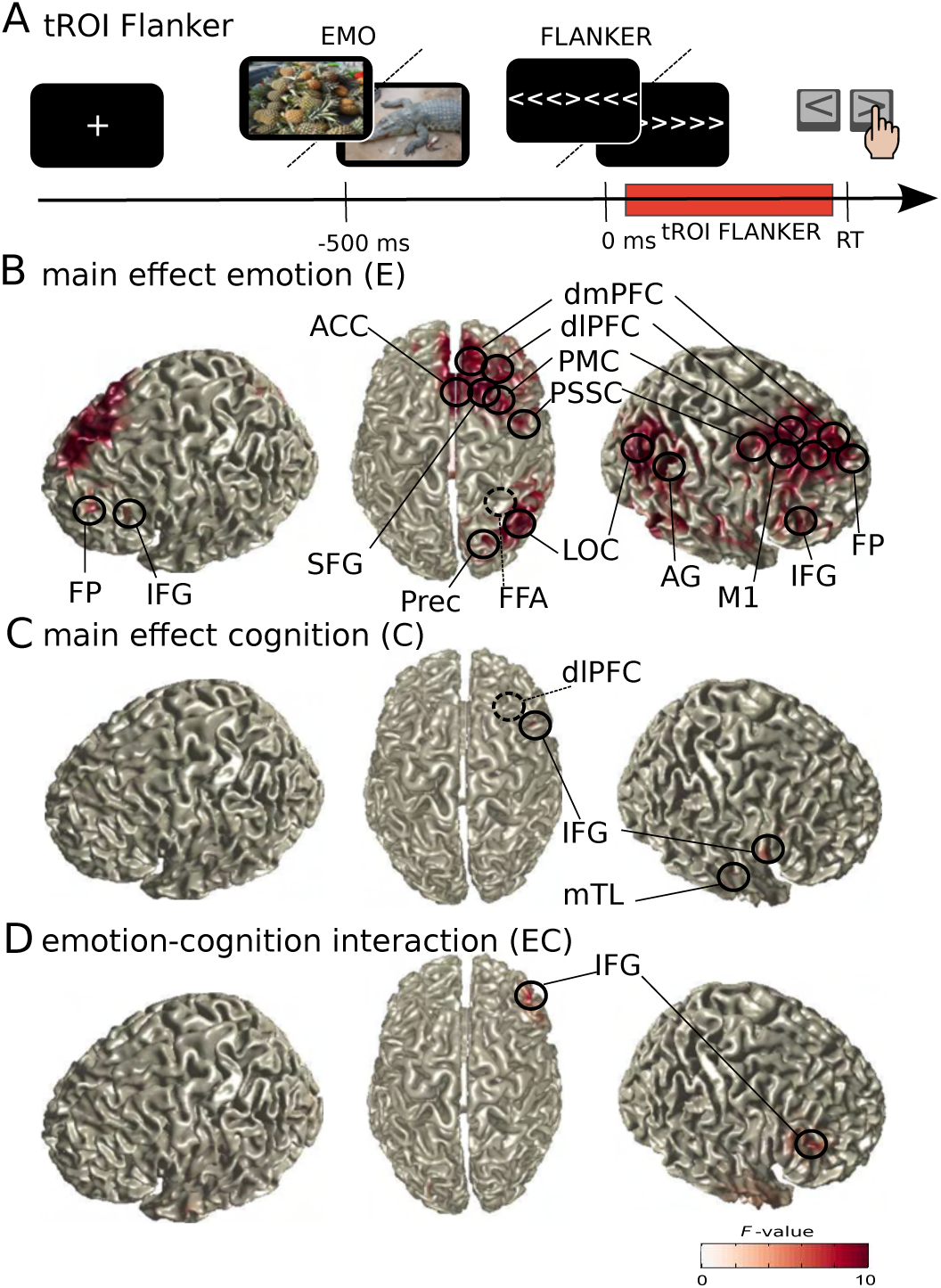
Source activity in the *β*-band (9-33 Hz). Surface plots show significant F-values of ANOVA (cluster-based permutation test, Bonferroni corrected for four frequency bands and two time windows, alpha and clusteralpha *<*0.00625, n=103). Peak voxels (local extrema) of significant clusters are indicated with circles and labels (for MNI coordinates, see 1, for a detailed description of the methods, see Source analysis and Statistical analyses of source reconstruction results). Dashed circles mark peak voxels invisible in this representation. A, Task design and analyzed tROI; B, main effect of emotion (E); C, main effect of cognition (C); D, interaction effect between emotion and cognition (EC). Anatomical regions: ACC, anterior cingulate cortex; AG, angular gyrus; dlPFC, dorsolateral prefrontal cortex; dmPFC, dorsomedial prefrontal cortex; FFA, fusiform face area; FP, frontal pole; IFG, inferior frontal gyrus; LOC, lateral occipital cortex; M1, primary motor cortex; mTL, medial temporal lobe, PMC, premotor cortex; Prec, Precuneus; PSSC, primary somatosensory cortex; SFG, superior fusiform gyrus.

**Table 1.** Significant *β*-band sources during tROI Flanker. All significant sources in the *β*-band are given for the contrasts (Contr.) of emotion (E), cognition (C) and statistical emotion-cognition interaction (EC). The table lists sources (Region (label)) with respective coordinates and F-values. Furthermore, it shows the point estimate in terms of source activity (mean differences of first level statistics (i.e., t-statistics) for main effects, and mean double differences for interaction effects) and the bootstrap Confidence-Interval (95% CI). Additionally, Cluster F- and P-values are reported.

| Contr. | Region (label) | Coordinat.<br>(x,y,z) | Source<br>F-value | point estimate<br>(Mean) | Bootstrap<br>95% CI | Cluster<br>F-value | Cluster<br>P-value |
| --- | --- | --- | --- | --- | --- | --- | --- |
| E | Fusiform Face Area R (rFFA) | 35 -50 10 | 16.1438 | 0.4319 | [0.2187; 0.6455] | 5.2295e<sup>+0.3</sup> | <1.9996e-04 |
| E | dorsomedial Prefrontal Cortex R (rdmPFC) | 5 40 40 | 15.9367 | 0.4302 | [0.2212; 0.6483] | 5.2295e+0.3 | <1.9996e-04 |
| E | dorsolateral Prefrontal Cortex R (rdlPFC) | 25 30 60 | 13.5796 | 0.3963 | [0.1808; 0.6073] | 5.2295e+0.3 | <1.9996e-04 |
| E | lateral Occipital Cortex R (rLOC) | 35 -70 30 | 13.3564 | 0.3686 | [0.1680; 0.5689] | 5.2295e+0.3 | <1.9996e-04 |
| E | Frontal Pole R (rFP) | 5 60 40 | 12.8408 | 0.4005 | [0.1744; 0.6177] | 5.2295e+0.3 | <1.9996e-04 |
| E | Angular Gyrus R (rAG) | 45 -50 20 | 12.6571 | 0.3737 | [0.1714; 0.5859] | 5.2295e+0.3 | <1.9996e-04 |
| E | Superior Frontal Gyrus R (rSFG) | 25 10 60 | 12.4352 | 0.3686 | [0.1613; 0.5781] | 5.2295e+0.3 | <1.9996e-04 |
| E | Premotor Cortex R (rPM) | 45 0 50 | 11.2470 | 0.3553 | [0.1453; 0.5647] | 19.5099 | <1.9996e-04 |
| E | Inferior Frontal Gyrus R (rIFG) | 45 40 0 | 10.4409 | 0.3852 | [0.1510; 0.6221] | 5.2295e+0.3 | <1.9996e-04 |
| E | Precuneous Cortex R (rPrec) | 15 -70 50 | 9.7742 | 0.3459 | [0.1311; 0.5695] | 5.2295e+0.3 | <1.9996e-04 |
| E | Frontal Pole L (IFP) | -25 50 10 | 9.6548 | 0.3641 | [0.1238; 0.5947] | 25.9950 | <1.9996e-04 |
| E | Primary Somatosensory Cortex R (rPSSC) | 45 -10 30 | 8.8410 | 0.3268 | [0.1172; 0.5519] | 5.2295e+0.3 | <1.9996e-04 |
| E | Inferior Frontal Gyrus L (lIFG) | -35 30 10 | 8.7014 | 0.3131 | [0.1040; 0.5265] | 8.7014 | <1.9996e-04 |
| E | Primary Motor Cortex R (rPrMC) | 35 -20 50 | 8.2433 | 0.3131 | [0.1016; 0.5327] | 8.2433 | <1.9996e-04 |
| C | dorsolateral Prefrontal Cortex R (rdlPFC) | 45 20 30 | 8.5053 | 0.3048 | [0.1082; 0.5220] | 8.5053 | <1.9996e-04 |
| C | medial Temporal Lobe R (rmTL) | 65 -10 -10 | 8.3118 | 0.3278 | [0.1194; 0.5716] | 8.3118 | <1.9996e-04 |
| C | Inferior Frontal Gyrus R (rIFG) | 55 10 10 | 7.9618 | 0.3034 | [0.1010; 0.5278] | 7.9618 | <1.9996e-04 |
| EC | Inferior Frontal Gyrus R (rIFG) | 45 40 20 | 9.6267 | -0.5948 | [-0.9589; -0.1966] | 9.6267 | <1.9996e-04 |

### Source-level activity confirms a functional parcellation of rIFG

The inspection of the IFG sources showed that they were located in different anatomical subdivisions of the rIFG: in the anterior ventral *pars orbitalis* (IFG_Orb_) for the main effect of emotion, in the posterior *pars opercularis* (IFG_Op_) for the main effect of cognition, and in the anterior dorsal *pars triangularis* (IFG_Tri_) for the interaction effect of emotion and cognition. To confirm that activity of these three reconstructed sources indeed originated from three separate sources and not from one underlying source, we correlated activities in the virtual channels reconstructed at the peak coordinates obtained from the beamformer source reconstruction with each other. Pairwise correlations between IFG subdivisions existed (*r* = 0.2844 for IFG_Tri_ and IFG_Op_, *r* = 0.3600 for IFG_Tri_ and IFG_Orb_, and *r* = 0.2937 for IFG_Orb_ and IFG_Op_), but were far from unity, suggesting partially independent sources and functional segregation of emotion/cognition processing within rIFG.

Notably, these three EEG sources are in close correspondence with three of five rIFG clusters indicated by an fMRI meta-analysis [15] and are located within the respective clusters with striking anatomical closeness to their respective centers-of-gravity, i.e., a maximum distance below 10 mm (see Fig 3, A). In line with the model by Hartwigsen and colleagues [15], the IFG_Op_ source, revealed by our cognition contrast, is located in a region associated with task execution, compatible with an involvement in stimulus and response interference processing in our Flanker task. The IFG_Orb_ source, revealed by our emotion contrast, is located in a region associated with emotion or social processing, compatible with an involvement in emotional processing in our emotional Flanker task. Finally, the IFG_Tri_ source, revealed by our statistical emotion-cognition interaction contrast, is located in a more dorsal region associated with reasoning and adaptive control, compatible with interaction or integration of emotion and cognition in our emotional Flanker task.

**Fig 3.**
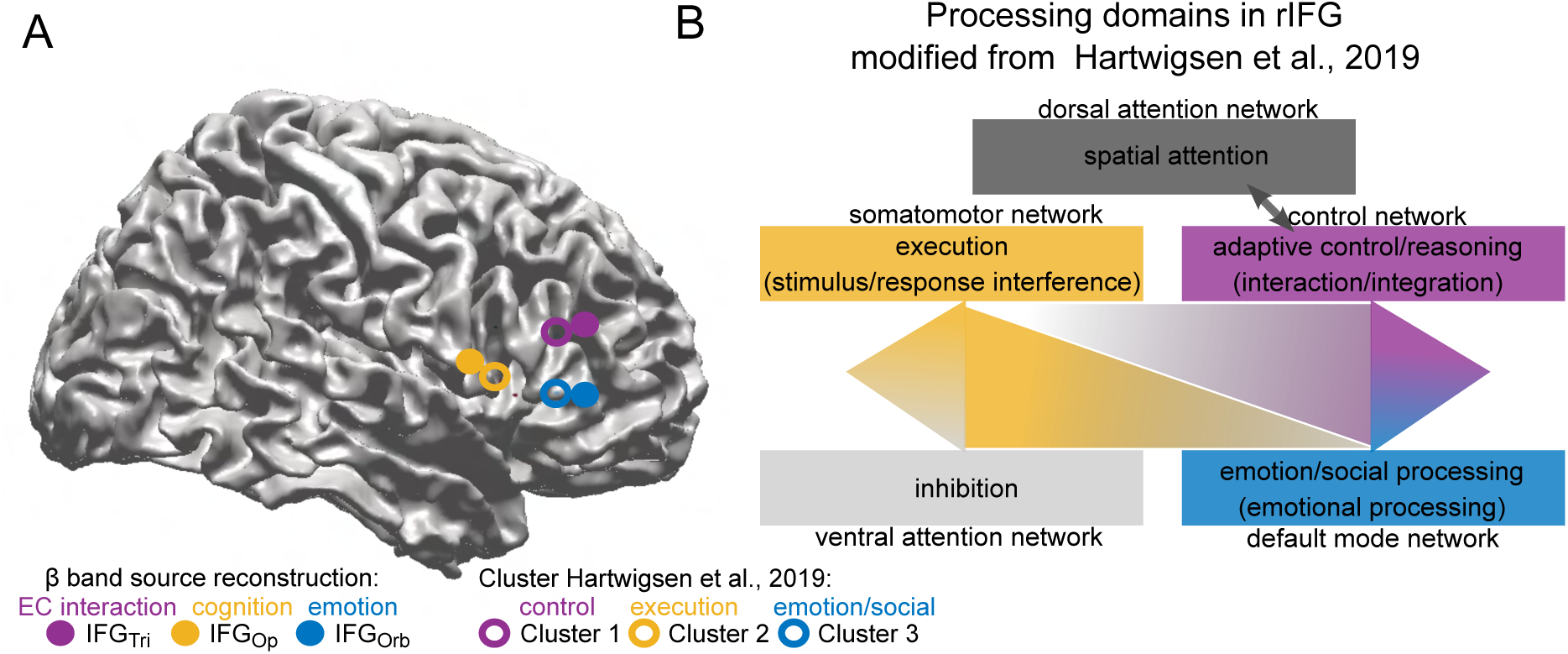
Segregation of task-related *β* sources within rIFG. A, Three separate *β* sources within rIFG (IFG_Orb_, IFG_Op_ and IFG_Tri_ represented by dots) were revealed by contrasts between task conditions (emotion: blue; cognition: yellow; statistical emotion-cognition interaction: purple), corresponding to anatomical subdivisions of IFG (*pars orbitalis*, *pars opercularis* and *pars triangularis*) and close to the centers of gravity of functional clusters (1-3, represented by circles; cluster 1: adaptive control/reasoning, purple; cluster 2: execution, yellow; cluster 3: emotion/social cognition, blue) obtained from fMRI meta analysis ([15]) B, Revision of the model by Hartwigsen and colleagues [15] indicating different processing domains in rIFG with incorporation of our findings (as indicated by brackets).

### Temporal Generalization analysis reveals an emotion-cognition interaction effect in IFG pars triangularis during early cognitive processing

In order to investigate the precise temporal dynamics of emotional versus cognitive processing, and of their interaction, we performed a temporal generalization classification in each rIFG subdivision with a support vector machine (SVM) (Fig 4) classifying the task conditions based on source-localized neural activity time courses. Thus, we were able to (1) classify between emotion (E), cognition (C), and the statistical emotion-cognition interaction (EC) processing, (2) determine the temporal unfolding of information processing in relation to these conditions, and (3) determine whether shared or isolated processing preferences exist in these rIFG subdivisions (see Fig 3, A). This analysis revealed that information related to emotion processing can be detected long after presentation of the emotional stimulus: we received a robust classification above chance level for emotion in all three IFG subdivisions starting around 200 ms before Flanker stimulus onset and extending until around 400 ms after Flanker stimulus presentation (p*<*0.0028, n=103, maximum classification accuracy and time point (Bootstrapping mean [95% CI]) was 60.1% [59.9; 60.2] at −18.5 ms [−20.8; −16.2] in IFG_Orb_, 59.5% [59.4; 59.7] at 2.6 ms [−1.3; 6.5] in IFG_Op_ and 58.5% [55.4; 58.7] at 56.0 ms [51.8; 60.2] in IFG_Tri_. The classification performance of the emotional process alternated between periods of high and low decoding accuracy, indicating that emotional information recurrently occupied processing resources. In contrast, accuracy of cognitive classification after Flanker signal onset continuously increased with time up to approximately 350 ms and remained relatively high until 500 ms post Flanker stimulus onset (p*<*0.0028, n=103, maximum classification accuracy and time point (Bootstrapping mean[95% CI]) was 64.9% [64.7; 65.1] at 354.1 ms [351.1; 357.1] in IFG_Orb_, 65.3% [65.1; 65.5] at 365.4 ms [362.5; 368.2] IFG_Op_ and 65.0% [64.8; 65.2] at 339.0 ms [189.8; 286.0] in IFG_Tri_ – indicating that resources devoted to this process ramp up (fast) over time. As expected, cognitive processes could reliably be classified above chance level after onset of the Flanker stimulus in all three subdivisions. Importantly, at the moment when cognitive information processing entered the network, i.e., when the SVM first classified the cognitive task conditions correctly, the strongest temporally specific statistical interaction effect between emotion and cognition took place in IFG_Tri_ (p*<*0.0028, n=103, maximum classification accuracy (mean, [95% Bootstrap-CI]) was 55.8% [55.3; 55.9] at 216.8 ms [214.5; 219.0]). Of note, the strongest interaction effect thus did not occur when the decodability of both processes was highest, but rather when the cognitive process entered the emotionally loaded network, presumably introducing a competition over processing resources. Together with our beamforming results, these findings strongly imply that IFG_Tri_ serves as the critical subdivision of IFG for emotion-cognition interaction.

**Fig 4.**
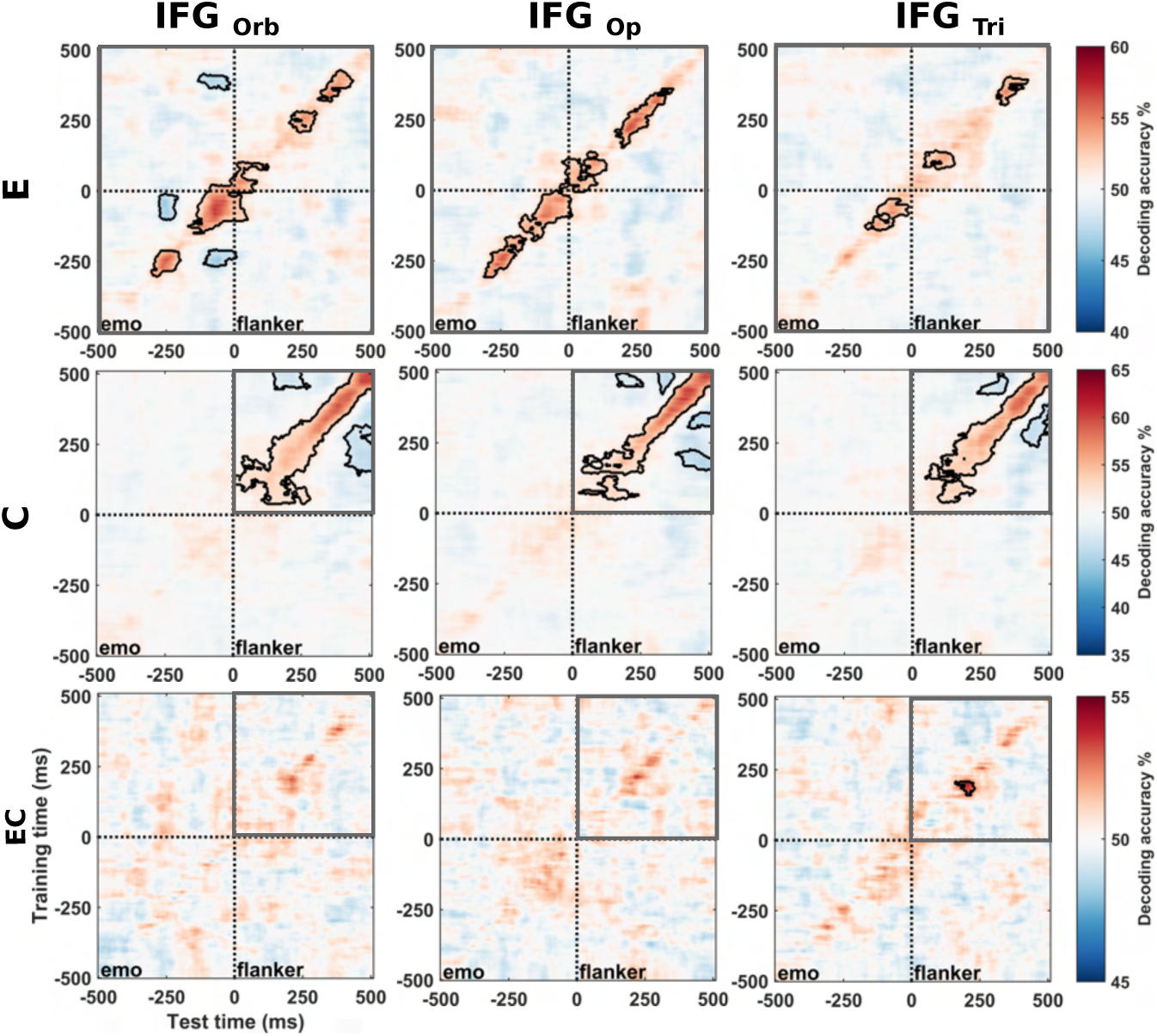
Temporal Generalization of task condition classification. Temporal generalization matrices for the three contrasts emotion (E), cognition (C) and their interaction (EC) (rows) in the three IFG subdivisions IFG_Orb_ (emotion), IFG_Op_ (cognition) and IFG_Tri_ (EC interaction) (columns). In each panel, a classifier was trained for each time sample (vertical axis: training time) and tested on all other time samples (horizontal axis: testing time), for each subject separately. The resulting individual decoding matrices were averaged across subjects, decoding contrasts and IFG subdivisions. Black contours indicate significant decoding values. Statistical tests used a cluster-size permutation procedure using a cluster defining threshold of p *<* 0.0028, and a corrected significance level of p *<* 0.0028 (n = 103). Grey boxes indicate time window of statistical test: contrast E was computed over the full time-range (−0.5-0.5 ms); since effects of contrasts C or EC can logically be present only after Flanker stimulus onset, a restricted time-range of 0-0.5 ms was used for these contrasts. Dotted lines indicate Flanker stimulus onset (0 (ms),”Flanker”).

### Integration and segregation of emotional and cognitive information within IFG

Given the functional segregation of IFG subdivisions found above and in [15] and the specific and behaviorally relevant role of IFG_Tri_ for emotion-cognition interaction, as well as the known anatomical interconnectivity of IFG subdivisions [16], we next aimed to understand the exact temporal and spectral dynamics of information flow within rIFG. To this end, we computed spectrally-resolved conditional Granger Causality (cGC) in two time windows (i.e., an early processing (visual to cognitive) phase [50-300 ms] and a late processing (cognitive to motor) phase [250-500 ms]) after Flanker stimulus onset between the three IFG subdivisions obtained from our source reconstruction contrasts cognition (IFG_Op_), emotion (IFG_Orb_) and statistical emotion-cognition interaction (IFG_Tri_)(see Non-parametric Granger Causality Analysis and Statistical testing of cGC for detailed description). Results showed significant spectral-cGC differences between IFG subdivisions, with the early time window (50-300 ms) primarily showing main effects of emotion in the *β* frequency (18-24 Hz, *p <* 9.99*e* − 05, *n* = 103, see also 2 for an overview), indicating lingering emotional processing. During this time window, IFG_Orb_ and IFG_Tri_ acted as both senders and receivers of emotional information, while IFG_Op_ only served as a receiver. In contrast, cognitive information was only sent from IFG_Orb_ to IFG_Tri_, at a frequency of 10 Hz (*p <* 9.99*e* − 05, *n* = 103) (Fig 5, Table 2).

**Fig 5.**
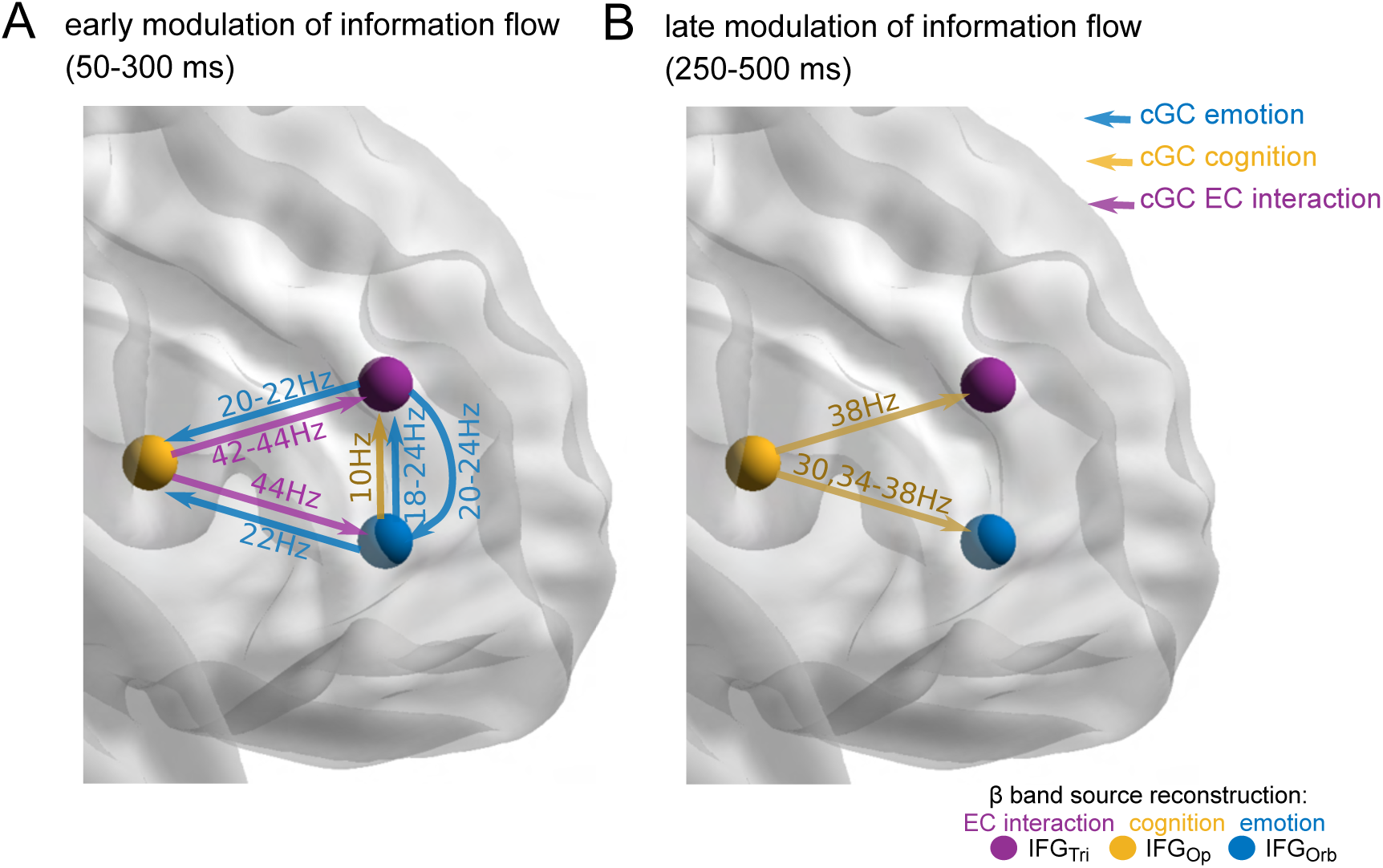
Spectrally-resolved conditional Granger Causality (cGC) within IFG subdivisions. Blue (IFG_Orb_, emotion (E)), yellow (IFG_Op_, cognition (C)), and purple (IFG_Tri_, statistical emotion-cognition interaction (EC)) nodes refer to the IFG sources in the *β*-band (see Fig 2 and Fig 3). Arrows indicate a significant spectral-cGC modulation (information flow) related to main effects of emotion (E, blue) and cognition (C, yellow) and their interaction (EC, purple) (see Methods), respectively. A cluster-based permutation ANOVA was used to identify significant differences between E, C, and EC contrasts. Significant differences after Bonferroni correction (*n* = 103 for all links, six links and two windows tested, *α_crit_* = 0.05*/*12) are reported with their significant spectral range. A, spectral cGC at early processing, 50-300 ms after Flanker stimulus onset. The modulation of information flowing between IFG nodes is mostly related to emotion and the statistical emotion-cognition interaction (blue and purple arrows). B, spectral cGC in the later time window 250-500 ms after Flanker stimulus onset. The modulation of information flow is mostly related to cognition (yellow arrow). Statistics were computed for n=103. For an overview of significant participants, links, and frequencies, please see 2.

**Table 2.** Overview of significant links within IFG. Table shows in which contrast, temporal region of interests (toi), frequencies (Hz) a link was significant in for how many participants (N(sig.)) with respective p-values of frequency-resolved group-level statistics. Point estimates (mean) and 95% confidence intervals from bootstrapping are reported. The frequency-agnostic individual level statistic was done for n=103; all reported links reached the lowest possible p-value in the permutation test. The threshold for significance was *p <* 0.0042. See Fig 5.

| Contrast | toi | Hz | Link | N(sig.) | p-values | p.e.<br>(Mean) | Bootstrap<br>95% CI |
| --- | --- | --- | --- | --- | --- | --- | --- |
| E | early | 22-22 Hz | IFG<sub>Orb</sub> → IFG<sub>Op</sub> | 103 | 9.999e-0.5 | 0.0059 | [0.0022; 0.0098] |
| E | early | 18-24 Hz | IFG<sub>Orb</sub> → IFG<sub>Tri</sub> | 103 | 9.999e-0.5 | -8.8583e-04 | [-0.0049; 0.0029] |
| E | early | 20-22 Hz | IFG<sub>Tri</sub> → IFG<sub>Op</sub> | 103 | 9.999e-0.5 | 0.0072 | [0.0033; 0.0118] |
| E | early | 20-24 Hz | IFG<sub>Tri</sub> → IFG<sub>Orb</sub> | 103 | 9.999e-0.5 | 0.0083 | [0.0041; 0.0125] |
| C | early | 10-10 Hz | IFG<sub>Orb</sub> → IFG<sub>Tri</sub> | 103 | 9.999e-0.5 | -0.0077 | [-0.0127; -0.0025] |
| EC | early | 42-44 Hz | IFG<sub>Op</sub> → IFG<sub>Tri</sub> | 103 | 9.999e-0.5 | 0.0135 | [0.0075; 0.0229] |
| EC | early | 44-44 Hz | IFG<sub>Op</sub> → IFG<sub>Orb</sub> | 103 | 9.999e-0.5 | 0.0118 | [0.0061; 0.0204] |
| C | late | 30-30 Hz | IFG<sub>Op</sub> → IFG<sub>Orb</sub> | 103 | 9.999e-0.5 | -0.0044 | [-0.0076; -0.0017] |
| C | late | 34-38 Hz | IFG<sub>Op</sub> → IFG<sub>Orb</sub> | 103 | 9.999e-0.5 | -0.0051 | [-0.0083; -0.0025] |
| C | late | 38-42 Hz | IFG<sub>Op</sub> → IFG<sub>Tri</sub> | 103 | 9.999e-0.5 | -0.0053 | [-0.0088; -0.0026] |

While receiving emotional information in the *β* frequency band, IFG_Op_ sent information related to statistical emotion-cognition interaction to both other subdivisions, at a frequency in the low *γ* range (42-44 Hz, *p <* 9.99*e* − 05, *n* = 103). Thus, emotion-cognition interaction seems to take place when the network is loaded with emotional processing, but before cognitive processing is firmly established. This finding is fully in line with findings from our temporal generalization analysis (Fig 4). During the late time window (250-500 ms) a shift in processing style within the IFG from emotional to cognitive processing occurred, with cognitive information flowing primarily from IFG_Op_ to both other IFG subdivisions in the high *β*/low *γ* frequency range (30 to 38 Hz, *p <* 9.99*e* − 05, *n* = 103). Of note, these frequency ranges are similar to the ones reported by Schaum and colleagues [11] in relation to a purely cognitive response inhibition task. In summary, while information related to emotional processing and emotion-cognition interaction, is distributed during an early processing phase after Flanker stimulus onset in the IFG network, information related to cognitive processing is mainly sent by IFG_Op_ during a later phase. In addition, data clearly support functional integration of information in a temporal and frequency-specific manner in the IFG network concurrent with functional segregation of anatomical IFG subdivisions, as described above (Temporal Generalization of condition classification).

### Top-down long-range inter-regional modulation by IFG

Beyond the behavioral significance of IFG in emotion-cognition interaction, we next sought to understand whether IFG exerts top-down control [17] over other areas relevant for emotion-cognition interaction with known anatomical as well as task-relevant connections to IFG [16] – and could thus potentially serve emotion interference *inhibition*. As rIFG was the only source exhibiting a statistical emotion-cognition interaction effect in the *β*-band (see Fig 2), we therefore identified potential target sources of top-down control in other frequency bands. In the high-*γ* band we indeed found two posterior sources, i.e., Precuneus (, n=103, MNI peak coordinates =-5, y=-60, z=30, F-Value=7.8551, p*<*1.9996e-04) and visual area V2 (n=103, MNI peak coordinates =5, y=-80, z=20, F-Value=7.9948, p*<*1.9996e-04), that showed a statistical emotion-cognition interaction effect (see Supplementary results of source localization). Next, we thus conducted a spectrally-resolved cGC analysis including all three IFG subdivisions (IFG_Orb_, IFG_Op_, and IFG_Tri_) and both posterior sources (Precuneus and V2) (Fig 6) to reveal potential top-down modulations (see Non-parametric Granger Causality Analysis and Statistical testing of cGC for detailed description of cGC analysis). In the early time window [50-300ms] we found significant emotion effects on information flow (p*<*0.0042, Bonferroni corrected for time windows and links; details in Table 3) from IFG_Tri_ to Precuneus at 10 Hz (*p* = 0.00369, *n* = 97). In the later time window [250-500 ms] we found significant cognitive effects on information flow from IFG_Tri_ to Precuneus and V2 at 34-36 Hz (*p* = 0.00439, *n* = 97), and from IFG_Orb_ to Precuneus at 10-14 (*p* = 0.00309, *n* = 97) and 32-36 Hz (*p* = 0.00119, *n* = 97) and to V2 at 34-36 Hz (*p* = 0.00349, *n* = 85). Further, we found an emotion effect on information flow from IFG_Op_ to V2 at 30-32 Hz (*p* = 0.00559, *n* = 100) that approached significance. No significant interaction effect of emotion and cognition on information flow was found in the top-down direction from IFG to the posterior areas. However, we again observed a switch in processing style, as cGC from IFG_Tri_ signals emotional information in the early phase of processing, while in the later phase cGC from IFG_Orb_ signals cognitive information. These findings highlight a temporally-, spectrally- and context-specific long-range modulation from rIFG to Precuneus and V2.

**Fig 6.**
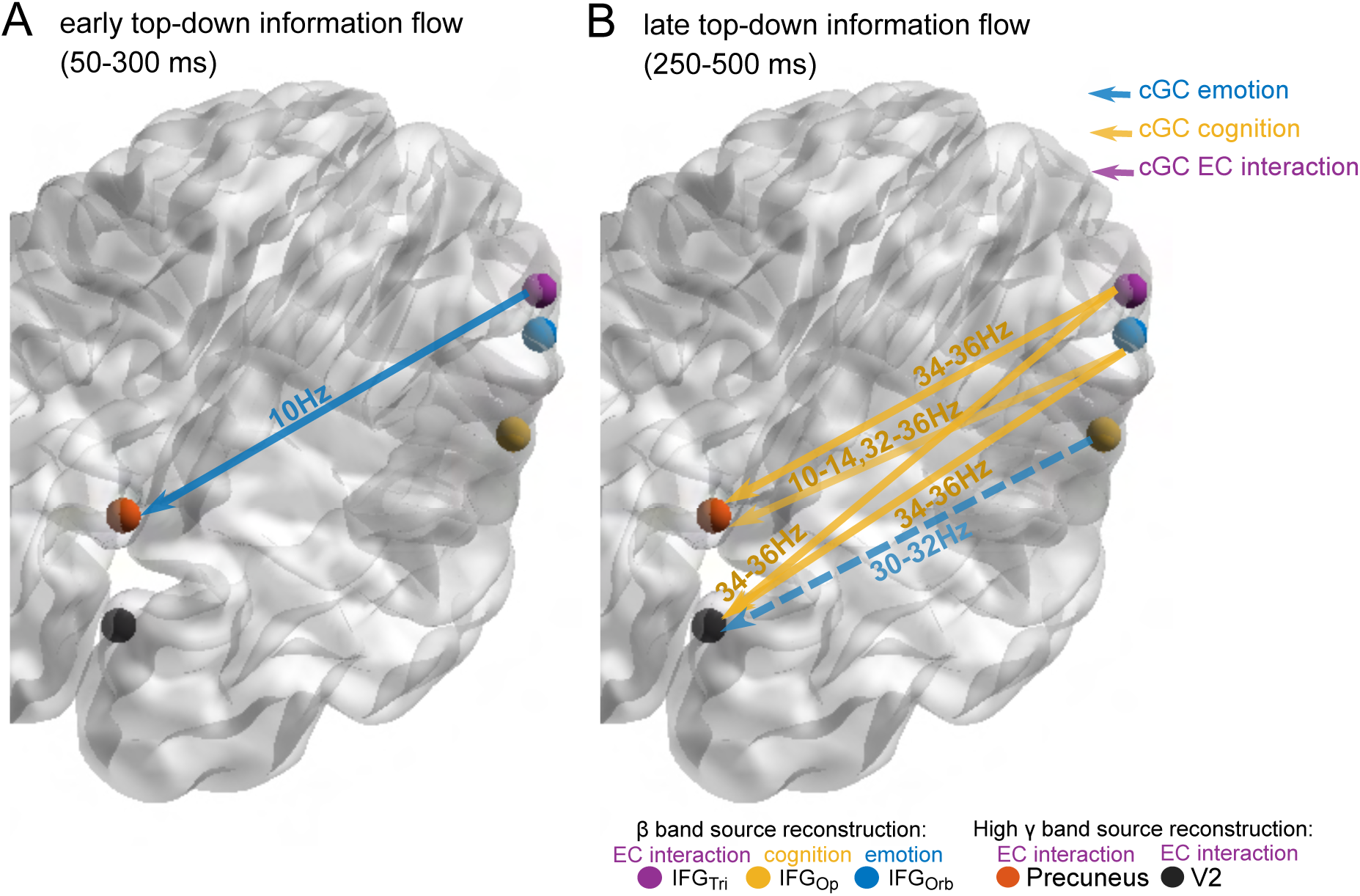
Spectrally-resolved conditional Granger Causality (cGC) between IFG subdivisions and Precuneus and V2. Blue, yellow and purple nodes refer to the IFG subdivisions reconstructed in the *β*-band (see Fig 2, Fig 3 and Fig 5); orange and black nodes refer to Precuneus and V2, reconstructed in the high *γ* band (see supplementary material). Blue, yellow and purple arrows indicate a significant spectral cGC modulation in emotion (E), cognition (C) and statistical emotion-cognition interaction (EC), respectively (see Methods for the definitions of these contrasts). Significant differences after Bonferroni correction (i.e., six links and two windows tested, p*<*0.0042) are reported as solid arrows with corresponding contrasts and significant spectral ranges. Close to significant links (p*<*0.00559) are denoted as dashed arrows. A, Spectral cGC at 50-300 ms after Flanker stimulus onset, showing IFG_Tri_-Precuneus modulation related to emotional processing. B, Spectral cGC at 250-500 ms relative to Flanker stimulus onset, showing IFG_Tri_ and IFG_Orb_ modulation of Precuneus and V2 related to cognitive processing. The frequency-agnostic individual level statistic was done for n=103. The frequency-resolved group-level statistics was done on significant participants, links, and frequencies, please see 2.

**Table 3.**
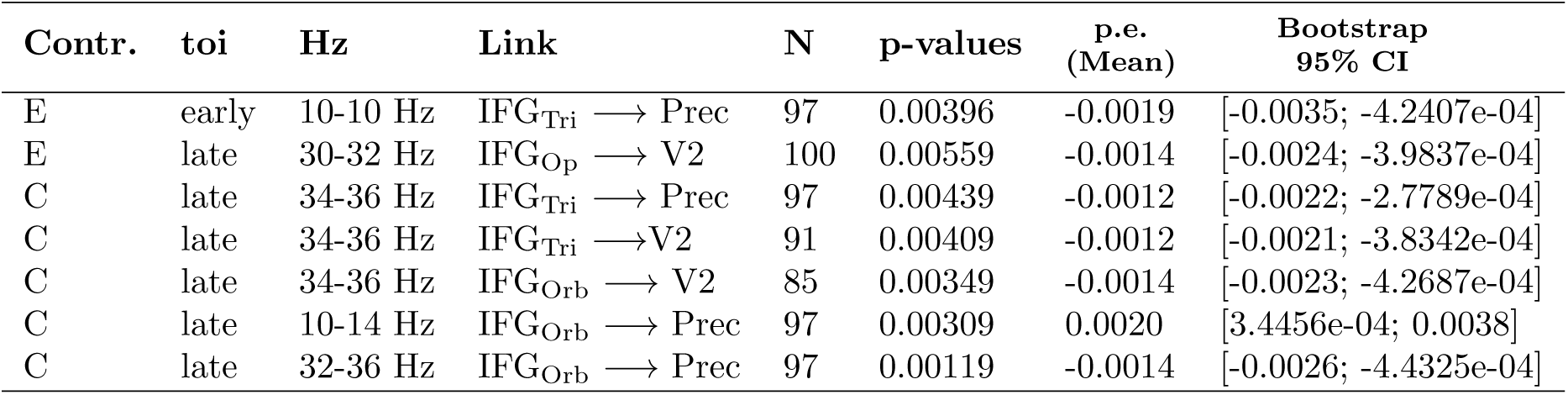
Overview of significant top-down links from IFG to Prec and V2. Table shows in which contrast (Contr.), temporal region of interests (toi), frequencies (Hz) the link was significant in how many participants (N) with respective p-values of frequency-resolved group-level statistics. Point estimates (mean) and 95% confidence intervals from bootstrapping are reported. The frequency-agnostic individual level statistic was done for n=103. Threshold for significance was *p <* 0.0042. See Fig 6.

| Contr. | toi | Hz | Link | N | p-values | p.e.<br>(Mean) | Bootstrap<br>95% CI |
| --- | --- | --- | --- | --- | --- | --- | --- |
| E | early | 10-10 Hz | IFG<sub>Tri</sub> → Prec | 97 | 0.00396 | -0.0019 | [-0.0035; -4.2407e-04] |
| E | late | 30-32 Hz | IFG<sub>Op</sub> → V2 | 100 | 0.00559 | -0.0014 | [-0.0024; -3.9837e-04] |
| C | late | 34-36 Hz | IFG<sub>Tri</sub> → Prec | 97 | 0.00439 | -0.0012 | [-0.0022; -2.7789e-04] |
| C | late | 34-36 Hz | IFG<sub>Tri</sub> → V2 | 91 | 0.00409 | -0.0012 | [-0.0021; -3.8342e-04] |
| C | late | 34-36 Hz | IFG<sub>Orb</sub> → V2 | 85 | 0.00349 | -0.0014 | [-0.0023; -4.2687e-04] |
| C | late | 10-14 Hz | IFG<sub>Orb</sub> → Prec | 97 | 0.00309 | 0.0020 | [3.4456e-04; 0.0038] |
| C | late | 32-36 Hz | IFG<sub>Orb</sub> → Prec | 97 | 0.00119 | -0.0014 | [-0.0026; -4.4325e-04] |

### The Top-down Influence from IFG to Precuneus and V2 explains the behavioral interference effect

Finally, we studied the behavioral relevance of the long-range inter-regional modulation by rIFG. To this end, we first determined which of the above identified combinations of links and frequencies were relevant for behavioral performance by correlating information flow, i.e., cGC, on these links with the behavioral measures in our emotional Flanker task, i.e., reaction time (stimulus interference inhibition) and response accuracy (response interference inhibition). We found that the change in information flow from IFG_Orb_ to Precuneus and V2, and from IFG_Op_ to V2 predicted behavior: For links and frequency combinations modulated by the factor emotion, cGC from IFG_Orb_ to V2 (34-36Hz connection; posterior median: *β* = −0.24, posterior HDI: [−0.67, 0.19]) was negatively correlated with reaction time and the same held for the cGC to Precuneus (32-36 Hz connection; posterior median: *β* = 0.58, posterior HDI: [0.16, 0.99] and 10-14 Hz connection; posterior median: *β* = −0.3, posterior HDI: [−0.55, 0.073]), which was also negatively correlated to response accuracy (32-36 Hz connection; posterior median: *β* = −0.29, posterior HDI: [−0.72, 0.13]). For links and frequency combinations modulated by the factor cognition, cGC (incongruent-congruent) from IFG_Op_ to V2 (30-32 Hz connection) was positively correlated with reaction time (posterior median:*β* = 0.13, posterior HDI: [−0.071, 0.32]) and negatively correlated with response accuracy (posterior median:*β* = −0.14, posterior HDI: [−0.34, 0.068]) (see Fig 7,A). All other combinations of links and frequencies were not informative for predicting the behavior.

**Fig 7.**
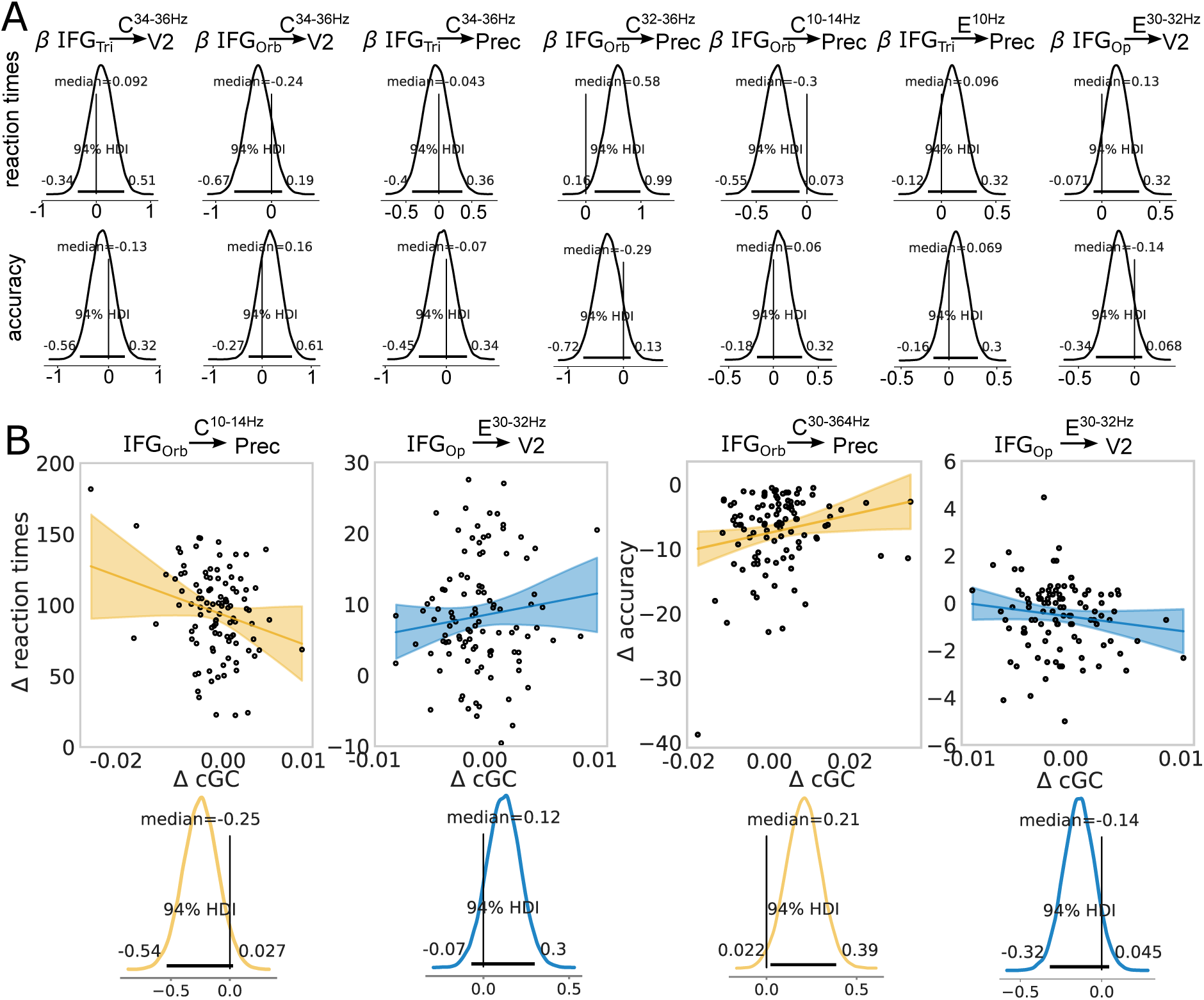
Spectrally-resolved cGC from IFG to Precuneus and V2 correlates with behavior. A, Posterior distributions of the Bayesian linear regression model parameters for models where cGC on various link and frequency combinations from IFG subdivisions to Precuneus (Prec) or visual area V2 (see black arrow above posterior) served as predictors, and behavioral measures (reaction time and accuracy) as outcomes. Contrast and frequency information can be found above the black arrows. Inset values indicate the median and 94% credible intervals of the posteriors; the latter values are also represented by the horizontal bars in black. Links with a 94% credible interval that were entirely positive or negative were used for subsequent Bayesian multiple linear regression models. B. Linear regression for a regression of the *differences* between conditions (E, C) in behavior (reaction times, accuracies) on the *differences* between conditions for cGC (ΔcGC), for cGC from IFG_Orb_ to Prec and V2, and IFG_Op_ to V2, respectively. Differences were encoded as, i.e., negative-neutral (diagrams with blue colors) and incongruent-congruent (diagrams with yellow colors). Corresponding marginal posterior distributions are shown below. See Correlation of top-down Granger Causality with behavior for a detailed description

**Fig 8.**
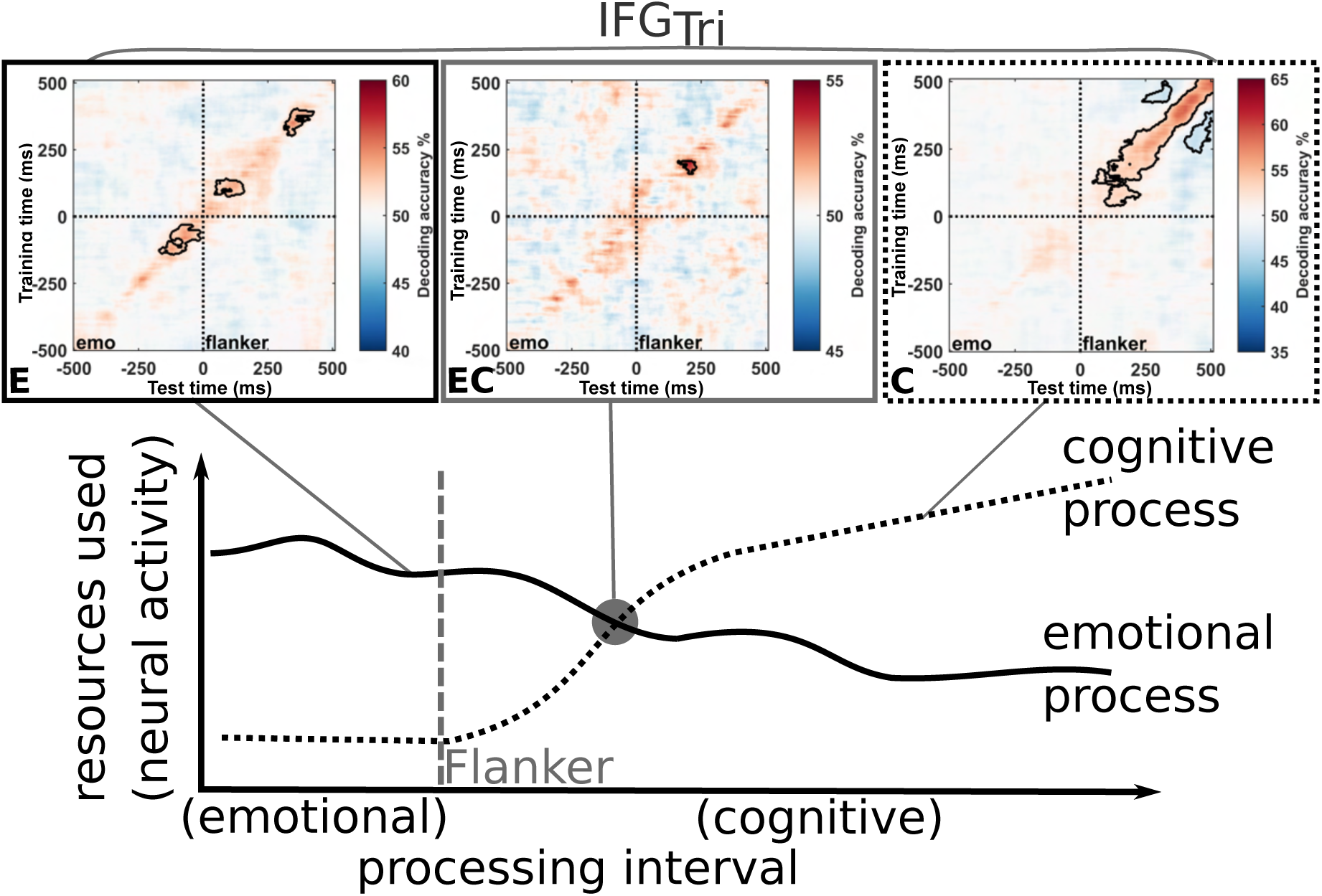
Neurophysiological model of emotion-cognition interaction. Emotional and cognitive processing compete for limited resources, with resources allocated to emotional processing decreasing during the ramp-up of cognitive processing. Emotion-cognition interaction occurs in rIFG*_tri_* at the cross-over of resource allocation to emotional and cognitive processing (also compare Fig 4).

Second, we computed Bayesian linear models to predict the interference effects (i.e., with Δ RT or Δ acc as dependent variables instead of RT and acc) caused by emotion and cognition, respectively, on behavioral performance, based on a model including all links originating in one subdivision as factors. In other words we correlated differences in information flows with differences in behavior: That is, the difference for the cognitive contrast in information flow from IFG_Orb_ to both Precuneus (32-36 Hz connection and 10-14 Hz connection) and to V2 (34-36 Hz) were used as factors in the same model to predict Δ reaction time (incongruent - congruent). Fig 7,B shows that the difference in top-down modulation (incongruent - congruent) from IFG_Orb_ to Precuneus was negatively associated with the cognitive interference effect on reaction time (10-14 Hz connection; posterior median:*β* = −0.25, posterior HDI: [−0.54, 0.027]), i.e., the stronger the difference in cognitive information flow was, the less impaired was the participants’ behavior due to the incongruent Flanker stimulus (i.e., cognitive interference). No association was seen for the difference in cognitive information flow from IFG_Orb_ to V2 (posterior median:*β* = −0.044, posterior HDI: [−0.32, 0.23]) and IFG_Orb_ to Precuneus in 32-36 Hz (posterior median:*β* = −0.077, posterior HDI: [−0.26, 0.11]) (see). In contrast, the difference in emotional top-down modulation (negative − neutral) from IFG_Op_ to V2 was associated with the emotional interference effect on reaction time (negative − neutral) (posterior median: *β* = 0.12, posterior HDI: = [−0.07, 0.3]) and accuracy (posterior median: *β* = −0.14, posterior HDI: [−0.32, 0.045]), i.e., the stronger the emotional information flow from IFG_Op_ to V2 in the negative condition was, the worse the behavioral performance was in the negative compared to the neutral conditions for both stimulus and response interference. The difference in cognitive information flow (incongruent − congruent) from IFG_Orb_ to Precuneus was positively associated with the cognitive interference effect (32-36 Hz connection; posterior median: *β* = 0.21, posterior HDI: = [0.0022, 0.39])

In sum, top-down effects on links modulated by emotional processing negatively impacted goal-directed, task-relevant behavior in our task, while those on links modulated by cognitive processing improved it.

## Discussion

In this study, we interrogated the neural mechanisms underlying emotional interference in cognitive processing, hypothesizing that competition for shared resources may cause the well-known behavioral effects of emotional stimuli on cognitive processing [1, 18]. Moreover, we asked whether such a competition would be resolved locally at the site of competition or via further brain areas executing control over the locus of competition. We found that processing of emotion and cognition, and their interaction, all coincided in the rIFG. Yet, there were subdivisions of the rIFG more related to cognitive processing (IFG *pars opercularis*) and emotion processing (IFG *pars orbitalis*), while there was one subdivision that was strongly implicated in processing both (IFG *pars triangularis*), and that showed an interaction. In this latter area, our analysis showed evidence consistent with direct competition for resources at the moment when cognitive processing entered the rIFG. This is consistent with a resolution of the conflict between emotion and cognitive processing by the competition itself, i.e., the need for resources serving cognitive processing partially displaces emotion processing. This notion is supported by the observation that emotional interference was stronger when cognitive load was less. This notion is further supported by the switch of internal communication within rIFG from emotion processing to cognitive processing over time. In addition, we observed that various subregions of rIFG exert long-range influences on parietal and visual cortices; these influences may mediate emotional stimulus interference detrimental to performance in the cognitive task, as well as cognitive stimulus interference inhibition which is necessary to perform the Flanker task.

### Large-N, FEM based EEG source analysis as a valuable tool for cognitive neuroscience

As detailed in the introduction, we aimed at improving the spatial resolution of EEG to enable a high resolution spatio-temporal mapping of neural activity underlying emotion cognition interaction. Considering that the sources we found in the three functionally segregated subdivisions of the rIFG agree very well with both, locations and functional descriptions from fMRI meta-analyses [15], we consider this goal achieved here, justifying the effort required for studies of this kind.

### A novel neural mechanism of emotion-cognition interaction

This improved spatial resolution in combination with the inherent temporal resolution of EEG then enabled us identify a previously unknown short-lasting interaction between emotion and cognitive processing in one particular subdivision of the rIFG, which we interpret as a direct and short lasting competition for resources – where task-relevant cognitive processing displaces task-irrelevant emotion processing from the rIFG. This view is supported by the fact that a higher cognitive load in the incongruent cognition seems to be more successful in displacing emotion processing, as indicated by the smaller behavioral penalty of the stimuli with negative emotional valence when cognitive load was high. Given the relatively low affective significance of the emotional stimuli used in our experimental setting, the rIFG with its three subdivisions may well be the place where the competition of resources for executive control and emotion processing takes place within a dual competition framework [3, 4].

### A dual role for *β*-band activity

At the neurophysiological level, the emotion-cognition interaction in the rIFG was visible, specifically in *β*-band activity. This observation is consistent with previous studies demonstrating the *β*-band to be important for *cognitive* control functions [13] such as behavioral inhibition (e.g., [11]) and extends to response inhibition in the Flanker-task used here. Our study adds the finding that indeed emotion processing also draws on these processing resources in the *β*-band such that a behaviorally relevant interaction effect in the rIFG occurred.

### Long range interactions originating in rIFG mediate effects of cognitive and emotional interference

We were also interested in potential neural mechanisms of emotion interference and emotion interference *inhibition*, where one brain area exerts behaviorally relevant control over another area. We found evidence for long range influences from IFG to V2 mediating emotion interference effects (Fig. 7, blue regression lines). Here, a stronger influence from IFG to V2 decreased behavioral performance. Thus, these links seem to mediate a behaviorally detrimental interference by the preceding, task-irrelevant emotional stimulus. We also found long range influences from IFG to Precuneus and V2 related to cognitive (Flanker) stimulus interference effects (Fig. 7, yellow regression lines). Since stronger influences from IFG to Precuneus and V2 resulted in improved behavioral performance, we interpret the function of these latter links as cognitive stimulus interference inhibition [19] [13].

### A central role for the rIFG in inhibitory control

While it was previously known that the rIFG plays a role in a specific subcomponent of inhibitory control [11, 20], i.e., behavioral inhibition [13] we here found that rIFG is also involved in two other inhibitory subcomponents, i.e., stimulus interference inhibition (reaction time) and response inhibition (accuracy) as measured by the Flanker task. This suggests that rIFG plays a broader role in inhibitory control, not limited to specific inhibitory subcomponents. We also add further evidence for its role in emotion processing, which is not well studied and understood to date (see [15, 21, 22]).

Our findings, which link activity in rIFG to, both, emotional and cognitive processing (i.e., emotional and cognitive interference) as well as their interaction (i.e., emotional interference inhibition), support the idea that the rIFG plays a critical general role in interference inhibition, regardless of the type of information causing the interference. Such a central role in interference inhibition is even further supported by our observation of the long-range modulation that rIFG exerts over visual and parietal areas, explaining the behavioral emotional and cognitive interference effect.

Such a broad involvement of the rIFG in various processes requires large-scale interactions in the brain. In line with this notion, research by [23] suggested that the macaque Brodmann area 45, which corresponds to the rIFG *pars triangularis*, is part of a tightly integrated large-scale innermost core network that connects various brain regions, including premotor, prefrontal, occipital, temporal, parietal and thalamic regions, as well as the basal ganglia, cingulate cortex, and insula. These connections offer both direct and indirect pathways that could explain the observed long-range top-down effects of rIFG on V2 and Precuneus, which is in line with a recent study by Boen and colleagues [16]. The IFG as part of the prefrontal cortex has recently been suggested to also be microstructurally exceptionally well-suited to handle broad information integration and receive input from different regions [24].

### The rIFG as a key region in emotion cognition interaction and emotion interference inhibition

In sum, our interpretation is that on the neural level, the rIFG plays a critical role in solving resource competition between emotion and cognition by allocating resources to either process and thereby implements a mechanism consistent with emotional interference inhibition. The outcome of this process is then also transmitted to visual and parietal cortices via behaviorally relevant top-down modulations of V2 and Precuneus. These long-range modulations could bias visual sensory processing in favor of task-beneficial processing by reducing emotional or cognitive (stimulus) interference. This interpretation is supported by previous research indicating IFG’s involvement in visual processing, such as salience [25] or change detection [26], and its potential to alter sensory processing [17, 27]. Since the IFG is also part of the attentional network, such long-range modulations and the sensory biases induced by them might involve attentional processes as well [28]. Our findings of frequency-specific top-down modulation are consistent with previous research, implicating *α*/*β*-band (see [29–32]) and (high) *γ*-band [33] activity in inter-regional connections, feedback, inhibitory control, attention [34] and emotional perceptual processing, respectively.

### Implications for a general theory of emotion-cognition interaction

Our findings fit within Pessoa’s dual competition model, where competition can arise at the perceptual and executive level [3]. Such competition due to emotion (or motivation, as he described) can both enhance and impair behavioral performance in cognitive tasks. Pessoa’s framework is based, inter alia, on the limited resources theory by Norman [35], which proposes that systems have limited processing resources and must allocate them to competing processes, selecting the most relevant processes for goal-directed behavior [4, 36]. Such a competition for resources by emotional and cognitive processing is located to the rIFG in our study.

Our findings also support the idea by Lavie and colleagues [37] that spare capacities are used for task-irrelevant processes if resources are not fully consumed. This is because in our study, emotion interference effects are more pronounced when cognitive load is low, indicating a strong influence of emotional stimuli when processing resources are not required in full by cognitive processing.

However, when considering the above idea of resource competition, this latter observation leads to the initially puzzling question of why interference is stronger when -seemingly- less processing resources are employed overall, and the limited capacities of a pool of shared resources [35] should be less likely to be exceeded. This puzzle may be resolved by the observation that emotional stimuli receive processing resources automatically and passively, even if irrelevant [2]. Thus, the processing of emotional stimuli may strain or exhaust processing resources in the rIFG ‘by default’. As a consequence, cognitive processing may compete more successfully with interfering emotional processing when cognitive load is indeed high enough to displace the processing of the irrelevant emotional stimuli. We consider this to be one plausible explanation for the pattern of behavioral results observed in our study.

On the neural level, Pessoa suggests that the ACC is key brain region mediating the interaction between emotion/motivation and executive functions [3]. In fact, the ACC is known for its involvement in conflict monitoring [38, 39], but also cognitive and emotional processes [40, 41]. However, we found that although the ACC was affected by the emotional interference, as it showed main effects of emotion during Flanker stimulus presentation (i.e., tROI Flanker), it did not show a significant emotion-cognition interaction effect, and therefore is not centrally involved in emotional interference inhibition. The rIFG, on the other hand, showed significant interaction effects and thus may play a more important role in emotional and cognitive processing and might serve as a common resource pool besides the ACC in our task.

Besides the ACC, Pessoa considers the amygdala to have a central role in his model based on its extensive connectivity. While the amygdala did not show significant effects at all in our study, this does not rule out this aspect of Pessoa’s model as the absence of amygdala-related findings may simply be due to the difficulty of reconstructing subcortical regions with EEG and FEM models, or due to our conservative statistics.

In sum, our findings support the dual competition model, but highlight the rIFG instead of the ACC as a central locus of the competition. Furthermore, this competition seems to be always present due to emotional stimuli, even if task irrelevant, receiving preferential processing and using a large fraction of available resources. Thus, resource competition may show large effects even in situations or tasks where an exhaustion of limited resources naively seems less likely due to low cognitive loads.

### Potential Clinical Implications

Impairment to control interfering impulses, whether cognitive or emotional, is a hallmark of various psychiatric disorders, including attention-deficit-hyperactivity disorder [42], autism [43], borderline personality disorder [44], major depression [45], obsessive-compulsive disorder [46], or substance use disorder [47]. Of note, recent research by [13] suggests that different subtypes of interference control deficits are operative in different psychiatric disorders, implying that these constructs may not be universally applicable. In contrast, findings from our current study show that both stimulus and response interference inhibition are affected by emotional interference. Further research is thus needed to elucidate the exact endophenotypes and pathophysiology of (selective) impairments of inhibitory control in patients with psychiatric disorders. Our study found evidence of the involvement of the rIFG, specifically the *pars triangularis*, in emotional interference inhibition. Previous research has linked aberrant neural activation patterns in IFG to anxiety and depression [48] and obsessive-compulsive disorder [49], and suggests that rIFG is a convergent site for response inhibition and state anger [50]. Likewise, reduced rIFG activity was reported in the context of impaired emotional interference inhibition in patients with major depressive disorder [51]. Taken together, these and our own findings highlight the critical role of rIFG in emotional interference inhibition and its potential as a target structure for the therapy of mental disorders. The temporal and frequency information provided by our current study could guide the development of targeted interventions, e.g., by non-invasive (electric or magnetic) brain stimulation [52, 53]. However, further research in patients with psychiatric disorders is clearly needed to elucidate the neural mechanisms of dysfunctional emotion-cognition interaction with similar spatio-temporal precision as in this study, thus potentially identifying neural targets for engaging and modulating rIFG and its subregions for treatment of impaired inhibitory control in these disorders. In addition to the importance for pathological conditions, our results will inform future preventive interventions for self-regulation and resilience fostering [54].

## Conclusion

In summary, our study provides compelling evidence for the importance of the rIFG, particularly the *pars triangularis*, in the interaction between emotion and cognition. By determining this interaction in space, time, frequency, and information transfer with high spatial and temporal resolution, we have shed new light on the neurophysiological mechanisms underlying emotional and cognitive processing, and the crucial role of rIFG in emotional interference inhibition. Overall, our study contributes to a more integrated understanding of the interplay between emotion and cognition, with clinical implications for psychiatric disorders such as depression and anxiety.

## Materials and Methods

### Ethics information

The study and all experimental protocols were approved by the local ethics committees of the Medical Board of Rhineland-Palatinate, Mainz, Germany, and Johann Wolfgang Goethe-University, Frankfurt, Germany, (ethical approval: 837.074.16(10393)) and all participants were financially compensated for study participation.

### Participants

A total of 121 healthy human subjects participated in this dual-center study after written informed consent (i.e., 59 and 62 subjects at study sites Frankfurt and Mainz, respectively). All participants were screened for magnetic resonance imaging (MRI) exclusion criteria, mental health status (Mini-International Neuropsychiatric Interview [55]) and handedness (Edinburgh Handedness Inventory [56]) prior to study inclusion. Four participants had to be excluded due to technical failures of the stimulus presentation during task performance. The data of the remaining 117 subjects were used for the behavioral analysis. Another thirteen participants were excluded from electrophysiological analysis due to missing or incomplete MRI data sets (three subjects due to panic attacks in the MR scanner which led to a premature abortion of the measurement, one subject due to pain during MRI measurement which led to a premature safety abort of the measurement, one subject did not fit into the MR scanner due to excessive abdominal girth, and eight subjects withdrew study consent before MRI measurements took place) and one subject had to be excluded due to external noise during EEG recording which could not be controlled for. Thus, all presented electrophysiological data are from analyses of the remaining 103 participants (65 females; mean age ± SD, 25 ± 6 years), who completed the study.

### Questionnaires

After study inclusion and screening for MRI exclusion criteria (see above), participants completed questionnaires on demographic information (Gender, date, and place of birth, family origin, psychological diseases within the family, family tree, marital status), Health and Lifestyle (physical and mental diseases, height and weight, blood pressure, medication, usage of internet, working conditions, income, family relationships, MRI compatibility) and their drug consumption (Fagerström for nicotine consumption and the AUDIT for alcohol consumption) (secutrial, www.secutrail.com). Further questionnaires were the Trier Inventory for the Assessment of Chronic Stress (TICS), the Short Form 36 Health Survey Questionnaire, the General Health Questionnaire (GHQ-28), the Life events checklist from LHC (adapted from [57]), the Cognitive Emotion Regulation Questionnaire (CERQ), the Positive and Negative Affect Schedule (PANAS), the State-Trait Anger Expression Inventory (STAXI), the State-Trait Anxiety Inventory (STAI), the Barratt Impulsiveness Scale (BIS-11 [58]), the Behavioral Activation and Behavioral Inhibition Scales (BIS/BAS), the Edinburgh Handedness Inventory [56] and an Intelligence test (L-P-S Leistungsprüfsystem UT-3 [59]).

### Experimental Design

Data collection took place in Frankfurt (Site 1) and Mainz (Site 2), respectively. At Site 1, 59 participants and at Site 2, 62 participants took part in the experiment. We attempted to set up and standardize the procedures as closely as possible. The experiment comprised of three experimental days (Day 1-3), two EEG measurements (Day 1 and 2) and one fMRI measurement (Day 3). On Day 1, participants were screened for exclusion criteria (see subsection Participants). Additionally, on Day 1 and 2, participants filled out the study questionnaires (see subsection Questionnaires). In total, four different behavioral paradigms (emotional Stop signal Task, emotional Recent Probes Task, Cognitive Emotion Regulation Task and emotional Flanker Task) were performed by the participants on Day 1 and 2, with two tasks being performed per experimental day. For each task, the order of the tasks was counterbalanced within and across participants. During execution of the tasks, an electroencephalography (EEG), which was spatially localized using digitalized electrode localization and finite element head models (see Finite element head modeling (FEM) and Fig 9), was recorded simultaneously. In addition to task-related EEG, an eyes-closed resting state EEG (rsEEG, 10 min) was recorded, either on Day 1 or Day 2 prior to task-related recordings, with the order of rsEEG on Day 1 or Day 2 counterbalanced across subjects. On Day 3, a resting state fMRI as well as structural T1 and T2 MRI measurements were carried out on each participant. Here, only data from the emotional Flanker Task EEG recordings as well as structural MRI measurements will be reported (see Structural imaging and head modelling and Electroencephalography – recording and analysis).

**Fig 9.**
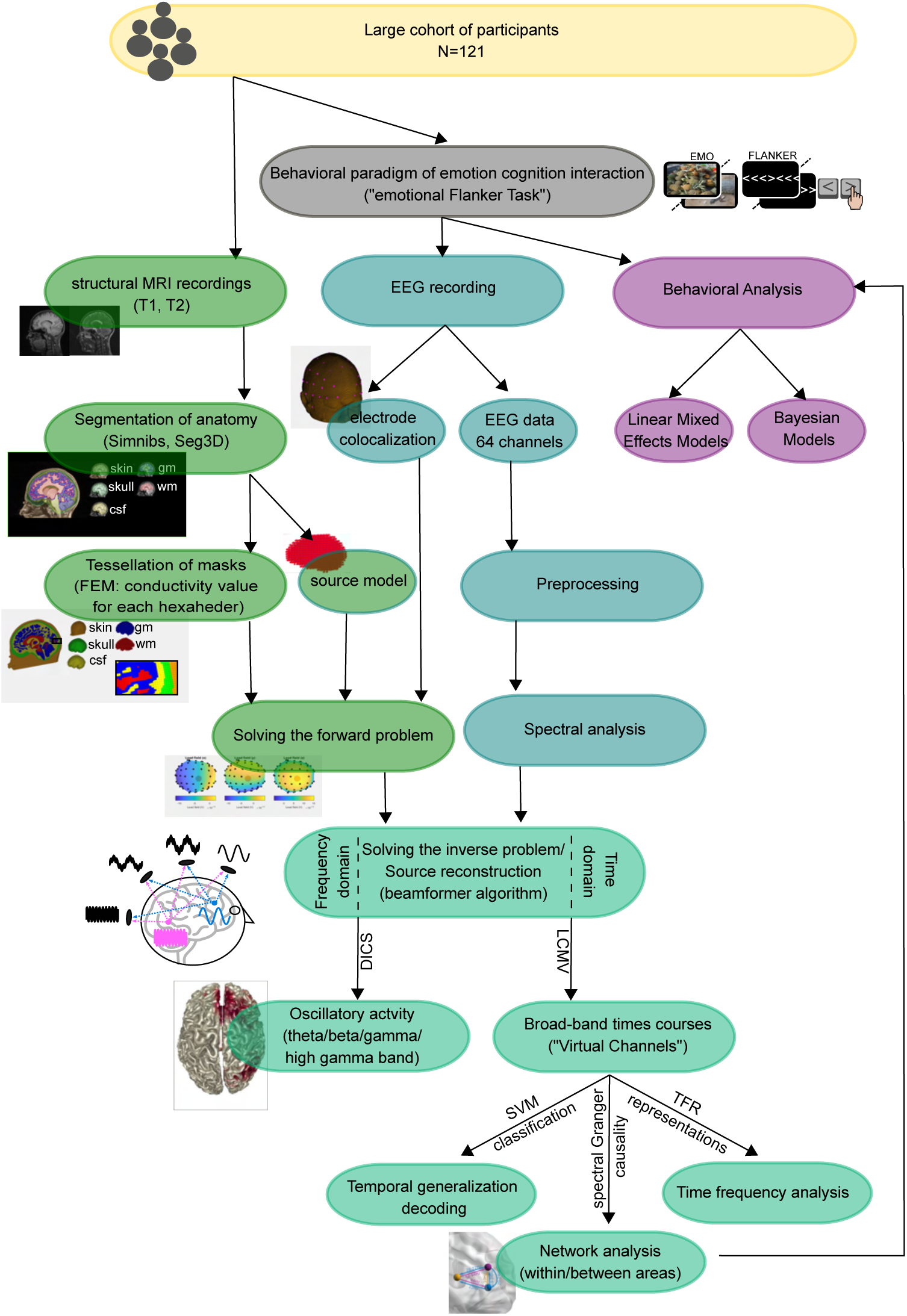
Pipeline for the analysis of emotion-cognition interaction. All major analysis steps are shown in the pipeline including EEG source reconstruction with Finite Element Headmodeling Finite element head modeling (FEM) based on digitalized electrode positions Electrode digitalization and spatial localization, structural analysis Structural imaging and head modelling and beamforming Source analysis, non-parametric statistics on source level Statistical analyses of source reconstruction results, temporal generalization of sources Temporal Generalization of condition classification, and Granger Causality analysis of information transfer between neural sources Integration and segregation of emotional and cognitive information within IFG, Non-parametric Granger Causality Analysis and behavioral analysis Bayesian analysis of behavioral data, Correlation of top-down Granger Causality with behavior.

### Task Design

For testing cognitive processing (i.e., stimulus interference and response selection interference operationalized by reaction time and accuracy, respectively, [13]) in the context of emotional distraction, a custom-made modified version of the Eriksen Flanker Task [60] was used, which was implemented in Presentation® (Version 18.1, Neurobehavioral Systems, Inc., Berkeley, CA, www.neurobs.com). Specifically, the Flanker stimulus in each trial was preceded by a picture with either neutral or negative emotional valence, taken from the International Affective Picture System (IAPS, [61]). Each trial consisted of (see Fig 1 A): fixation cross (central, on black background, duration: 1000 ms), emotional stimulus (IAPS picture, neutral and negative, duration: 500 ms) and Flanker stimulus (white arrows on black screen, with congruent or incongruent flanking arrows, duration: 1000 ms). The inter-trial interval (black screen) was 400 ms. Participants were instructed to indicate the direction of the central target arrow via left or right button presses (i.e., left and right ctrl buttons on regular keyboard) via the index finger of the respective hand and ignore the flanking arrows. In case of an incorrect (wrong button press) or a response that was too slow (reaction time *>* 1000 ms) an error/no response feedback was displayed on the screen for 300 ms. In total, 1120 trials in five blocks of 224 trials each (duration: 11 min) were performed during simultaneous EEG recording. IAPS pictures were used as emotional stimuli: 280 neutral pictures (mean valence was 5.15, mean arousal was 3.17) and 280 negative pictures (mean valence was 2.47, mean arousal was 6.41). Each emotional picture was repeated twice and with different Flanker stimulus. The combination of the emotional stimulus (negative/neutral) followed by the Flanker stimulus (incongruent/congruent) led to four different conditions, i.e., neural congruent: neutral picture followed by congruent Flanker, neutral incongruent: neutral picture followed by incongruent Flanker, negative congruent: negative picture followed by congruent Flanker, and negative incongruent: negative picture followed by incongruent Flanker. For each condition, 280 trials were presented with half of the trials showing a right- and left-side directed central target arrow, respectively. Blocks of trials were separated by 5 minutes to allow participants to relax in between blocks. To familiarize subjects with the task, prior to the recording, a training block with 20 trials with neutral pictures only was presented at the beginning of the experiment.

### Behavioral Response acquisition

Behavioral responses were analyzed in terms of reaction time and accuracy. Reaction time was measured as the time in ms after the Flanker stimulus onset until the button press of the subject. Only correct trials were considered for reaction time analysis. Reaction times faster than 90 ms were excluded, as reaction times in this range have been reported to be biologically implausible [62, 63]. Trials with a reaction time above 1000 ms were excluded too. Accuracy was determined as the ratio of all correct trials and all displayed trials of a given task condition, and calculated for each condition separately. The statistical analysis was performed on 117 subjects (see above).

### Bayesian analysis of behavioral data

The experimental effect of the factors emotion and cognition on reaction time and accuracy was determined with a Bayesian linear regression. We employed a hierarchical model and used a non-centered parameterization [64, 65] to foster convergence. We modelled the emotion and cognition effects and their interaction. For comparison to previous studies using frequentist statistics a similar mixed-effect model was also analyzed [66] with (RStudio Version 1.4.1106 © 2009-2021 RStudio, PBC); the corresponding mixed-effects modeling procedures and results are reported in the Supplementary information about behavioral effects of emotion-cognition interaction and. In the following, we describe the Bayesian model definition for reaction time and accuracy data. Single-trial values of reaction times were *z* -transformed and log-transformed (to account for their skewed distribution) before fitting the model, the estimated regression coefficients are reported with respect to these transformations. For the *i* th trial and the *j* th subject, we can define the likelihood of the reaction time given by *y_i,j_*as:

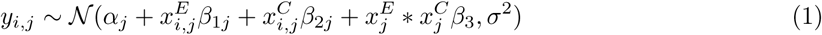

where for the *j* th subject, *α_j_* is the random intercept and encodes the grand average response time and *β*_1*j*_, *β*_2*j*_ are the random slopes which capture the differences in response times for emotion (negative vs. neutral), cognition (incongruent vs. congruent), respectively. Adding per-subject random slope parameters also for the interaction predictor term *β*_3_ did not improve the model and was thus left out. The term 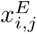 encodes the emotion condition (−0.5 = negative; 0.5 = neutral), the term 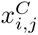 encodes the cognitive condition (−0.5 = incongruent; 0.5 = congruent). To obtain a more efficient sampling of the posterior, we inferred the random effects (intercepts and slopes) with a deterministic transformation of the parameters *α_j_* and *β*_1..2*j*_. More specifically, we did not infer individual slopes by modeling their values directly, for example saying that they are normally distributed around a group mean. Instead, we modelled their values relative to the mean (i.e., *β_offset_*) and allowed the effects to deviate from the mean to a certain degree (i.e., *σ_b_*).

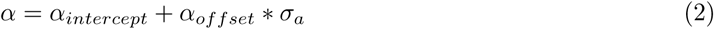

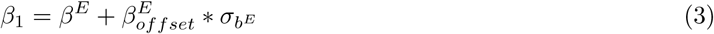

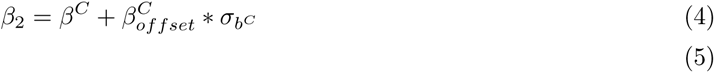

For the accuracy data we assumed *Y_i,j_* to have a Bernoulli distribution:

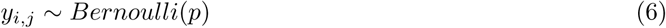

with probability of success *p_i_*, where in the logistic regression model the logit of the probability *p_i_* is a linear function of the predictor variable *x_i_*:

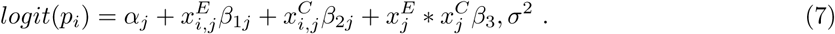

As above, we used the same non-centered parameterization. All parameters were assumed to be drawn from normal distributions. We used non-informative priors in both models that can be found in.

### Structural imaging and head modelling

#### Electrode digitalization and spatial localization

The position in space of all electrodes as well as the location of the nasion and fiducials were digitized and recorded with the ultrasonic sensor pen 3D-Digitizing System ELPOS and the respective ElGuide Software (ZEBRIS, Zebris Medizintechnik, Tübingen, Germany). Care was taken to ensure correct registration and spatial configuration of the electrodes. To this end, we created an average configuration file for each EEG cap size (56 cm, 58 cm, 60 cm) of all participants wearing the respective cap size. We then compared each electrode position of each individual electrode configuration to i) the respective electrode position of the average electrode configuration and ii) to the respective electrode position of the same participant’s electrode configuration at her/his second EEG session (not of interest for this study) via Euclidean distance. In case an electrode position exceeded 1 cm off reference, we replaced its position with the respective coordinates of either the average configuration or of the second EEG session configuration, depending on visual inspection to ensure correct electrode position.

#### Magnetic resonance imaging (MRI)

To create individual head models for source reconstruction, T1- and T2-weighted structural MRI datasets were acquired for each individual participant. To ensure later co-registration and alignment of the digitized electrode position, fiducials were used as anatomical markers. We added vitamin E pills at the fiducials positions when acquiring the MRI, using the same convention when marking them during the digitization of the electrodes’ location. At the study site 1 (Frankfurt), MRI data were acquired at a SIEMENS MAGNETOM Trio Syngo MR A35 3 Tesla MRI System (Erlangen, Germany) with an 8-channel head coil. A magnetization prepared rapid gradient echo (MPRAGE) T1-weighted sequence with fat suppression lasting 4:28 min (192 sagittal slices of 1 mm thickness with a distance factor of 50%, FOV 256 × 256 mm, 256 × 256 matrix, TR = 1900 ms, TE = 2.74 ms, flip angle = 9°, 1.0 × 1.0 × 1.0 mm3 voxels) and a turbo spin-echo sequence (TSE) sequence lasting 7:51 min (192 sagittal slices of 1 mm thickness with a distance factor of 0%, FOV 256 × 256 mm, 256 × 256 matrix, TR = 1500 ms, TE = 355 ms, flip angle = 0°, 1.0 × 1.0 × 1.0 mm3 voxels) were acquired. At study site 2 (Mainz), MRI data were acquired at a SIEMENS MAGNETOM TrioTim Syngo MR B17 3 Tesla MRI System (Erlangen, Germany) with a 32-channel head-coil. The specific parameters for each sequence were defined to match as closely as possible those from study site 1 given the respective specificities of the MRI system and the available software. That is, an MPRAGE T1-weighted sequence with fat suppression lasting 4:26 min (192 sagittal slices of 1 mm thickness with a distance factor of 50%, FOV 256 × 256 mm, 256 × 256 matrix, TR = 1900 ms, TE = 2.23 ms, flip angle = 9°, 1.0 × 1.0 × 1.0 mm3 voxels) and a TSE sequence lasting 7:50 min was used (192 sagittal slices of 1 mm thickness with a distance factor of 0%, FOV 256 × 256 mm, 256 × 256 matrix, TR = 1500 ms, TE = 339 ms, flip angle = 0°, 1.0 × 1.0 × 1.0 mm3 voxels) were used.

#### Finite element head modeling (FEM)

To create individual finite element head models, we used SimNIBS (Version 2.0.1, www.simnibs.org) with mri2mesh [67] and the FieldTrip-SimBio pipeline (Version 2.0.1) [10]. First, the mri2mesh pipeline from SimNIBS was used with the individual T1 and T2 images as input to create realistic individual segmentations. Hereby, T2-weighted images, in addition to T1-weighted images, are used to further improve the segmentation between skull and cerebrospinal fluid [68]. This pipeline involves Freesurfer [69, 70] and FSL [71] to automatically segment brain and non-brain tissue into five compartments (skin, skull, cerebrospinal fluid, white, and gray matter) [67]. The results of the automatic segmentations were visually inspected in Seg3D (version 2.2.1, Scientific Computing and Imaging Institute, University of Utah, https://www.sci.utah.edu/cibc-software/seg3d.html) to check for segmentation errors and manually correct them if necessary. The surface masks so generated were checked for holes and overlaps. Using the Fieldtrip-Simbio pipeline, the masks were then transformed into meshes consisting of hexahedrons, which subsequently were used to compute the individual FEM forward solutions. Following [72, 73] we assigned 0.33, 0.14, 1.79, 0.01 and 0.43 S/m as conductivity values for gray matter, white matter, cerebrospinal fluid, skull, and scalp. The FEM models were aligned to the digitized electrode positions via the nasion and fiducials. To reduce the space between the real recording site (lower part of the electrode) and the digitized site (upper part of the electrode), the electrode position of each electrode was projected down to the nearest point onto the skin. We warped a 3-D template grid with 10 mm spacing in Montreal Neurological Institute (MNI) from fieldtrip [74] to each individual head model covering the whole brain. Hereby, each grid point can directly be compared to the same grid points of other participants, as each point represents the same respective brain area. As a last step, we calculated the individual lead fields per warped grid point, representing the electrical field of the respective tissue, given the individual electrode position.

### Electroencephalography – recording and analysis

#### Data acquisition

Surface EEG was continuously recorded during the emotional Flanker task with an active 64-channel EEG system (actiCAP system, Brain Products GmbH, Gilching, Germany) using the standard 64Ch-actiCAP-Slim electrodes montage (Brain Products GmbH, Gilching, Germany). Four recording channels (predefined in the montage by brain products at position P7, P8, P9, P10) were placed near the outer right and left edges of the eye and 1 cm below and above the right eye to collect horizontal and vertical electrooculography (EOG), respectively. Therefore, EEG data was collected from only 60 channels. EEG data were recorded with BrainVision Recorder (Version 1.20.0601) with a sampling rate of 2.5 kHz, except for the first nine participants at Site 1, where a sampling rate of 5 kHz was used. Participants were placed centrally at about 80 cm distance in front of a standard PC monitor (ASUS VG278HV, 27”, 144Hz, 1920×1080, 1 ms GTG) within an EEG chamber with sound and RF shielding. In order to minimize artifacts during data acquisition, participants were instructed to relax their jaw and neck muscles and sit as still as possible during the recordings.

#### Preprocessing of EEG data

Preprocessing and analysis of the EEG data were performed with MATLAB R2012b (MathWorks Inc, Natick, MA, USA) and the FieldTrip toolbox (Version 20180729) [74]). First, data were low-pass filtered with a cut-off at 300 Hz. Subsequently, data were down-sampled to 500 Hz and cut into 1500 ms long epochs starting with the emotional picture onset. Other epochs were cut 2000 ms starting with the fixation cross onset and ending 500 ms after Flanker stimulus onset. Bad trials (i.e., with electrode jumps or muscle artifacts) were rejected using the FieldTrip automatic artifact rejection functions and line noise was filtered out with discrete Fourier transform filters at 50, 100, 150 and 200 Hz. Further artifacts (e.g., due to eye movements or heart beat) were removed using an extended infomax (runica) algorithm (provided in Fieldtrip/EEGLAB) [75]. Independent components were visually inspected and rejected based on topography and spectral power [76] and/or if there was a significant correlation (rho*>* 0.3) with either the EOG stimulus or with a prototypical “square-root” spectrum commonly observed in muscle artifacts [77]. Then data was re-referenced to the common average. This EEG data cleaning approach resulted in, on average, exclusion of 16.29 ± 29.72 (mean ± standard deviation) trials (range of excluded trials 0 to 177) out of a total of 1120 trials. To assure that statistical differences in subsequent comparisons were not caused by different numbers of trials per condition, we stratified the number of trials per condition per subject by randomly selecting for each condition the minimum amount of trials found across conditions in case of differences in the number of trials per condition. Furthermore, we only used correct trials, i.e., trials with correct button presses within required response times (*>* 90 ms, *<* 1000 ms).

#### Spectral analysis at sensor level

To define specific frequency bands for subsequent source reconstruction and further source-level analysis (see Source analysis), we first analyzed the spectral power at sensor level in order to find all potentially task relevant frequency bands. We obtained spectral power estimates between 3 and 120 Hz with Hanning tapers, over the tROI Flanker, and baseline period (see Temporal ROI definition), for all EEG channels and participants. In order to avoid circular reasoning and ‘double-dipping’ with respect to the subsequent statistical analysis comparing conditions in EEG source space, we performed sensor-level statistics by contrasting ‘pooled’ task conditions (all four conditions) with corresponding baseline periods ([78]). For this, we used a dependent samples t-test and a cluster-based correction method ([79]) to account for multiple comparisons across frequencies only (since data was averaged across channels and time). We employed a two-tailed test with an alpha threshold of *α* = 0.025 and cluster alpha of *α_cluster_*= 0.025. Finally, to delineate different frequency bands in the statistical analysis results, we identified the points of maximum curvature in the t-value vs. frequency curve by visual inspection. This allowed us to determine four non-overlapping frequency bands (a *θ* band from 5 to 9 Hz with a center frequency of 7 Hz (± 2 Hz spectral smoothing), a *β* band from 9 to 33 Hz with a center frequency of 21 Hz (± 12 Hz spectral smoothing), a *γ* band from 33 to 64 Hz with a center frequency of 49 Hz (± 16 Hz spectral smoothing) and a high *γ* band from 64 to 140Hz with a center frequency of 102 (± 38 Hz spectral smoothing) (see).

#### Temporal ROI definition

Two temporal regions of interest, one during the Flanker stimulus presentation (tROI Flanker) and another during the emotional picture presentation (tROI Emo) were defined (Fig 1) for each frequency band. To exclude early visual processing, the tROI Flanker started only 50 ms after the Flanker stimulus onset ([80, 81] and ended around the average minimum reaction time (430 ms) to exclude motor responses. This led to time windows of interest for each frequency band (beta, *γ*, and high *γ* band) with a duration of approximately 380 ms. The duration of tROIs varied slightly to match a full number of cycles of the center frequency of each frequency band, i.e., a center frequency of 21 Hz in the beta band resulted in a tROI of 380.95 ms (8 full cycles), a center frequency of 49 Hz in the *γ* band resulted in a tROI of 387.76 ms (19 full cycles) and a center frequency of 102 Hz in the high *γ* band resulted in a tROI of 382.35 ms (39 full cycles). As lower frequencies have longer wavelengths, we increased the time window for the *θ* band to obtain three full cycles of the center frequency of 7 Hz to 428.57 ms by starting at tROI FLANKER onset. The same procedure was applied to the tROI EMO during the emotional picture presentation. In the *θ* band, analogous to the tROI FLANKER, the tROI EMO was 428.57 ms long and started at the emotional picture onset; see Fig 1 B and Temporal ROI definition for definition of tROIs).

#### Source analysis

Beamformer source analysis was performed using the dynamic imaging of coherent sources (DICS) algorithm [82], with real-valued filter coefficients, as implemented in FieldTrip. This adaptive spatial filter estimates the power at each grid point based on individual lead fields (see Finite element head modeling (FEM)) and cross-spectral density matrices. The latter were computed for the two previously described ROIs (see Temporal ROI definition for further details) and for a baseline period (from −500 ms to −120 ms (and to −71.65 ms in the case of *θ*) before the onset of the emotional picture during the fixation cross) and four different bands. The frequency bands were defined as described in section Spectral analysis at sensor level, and consisted of a *θ* band with a center frequency of 7 Hz (± 2 Hz spectral smoothing), a *β* band with a center frequency of 21 Hz (± 12 Hz spectral smoothing), a *γ* band with a center frequency of 49 Hz (± 16 Hz spectral smoothing) and a high *γ* band with a center frequency of 102 (± 38 Hz spectral smoothing). Cross-spectral density matrix calculation was performed using the FieldTrip toolbox with the multitaper method [83]. We used a regularization of 5% ([84]). Beamformer filters were computed based on the data from both tROIs and the baseline across all conditions (’common filters’), thus assuring that differences in source activation do not arise from differences in filters derived from the individual conditions. Additionally, we report point estimates (mean) and 95% confidence intervals from bootstrapping with 5000 random resampling for each contrast: emotion (negative − neutral), cognition (incongruent − congruent) and emotion-cogniton interaction ((negative incongruent − neutral incongruent) − (negative congruent − neutral congruent)).

#### Statistical analyses of source reconstruction results

The source statistics comprised of two levels. First, we computed within-subject t-values by two-sided t-tests on single trial data for tROI Flanker versus baseline and for tROI Emo versus baseline (see Fig 1) for each task condition and grid point (dual state beamformer [85]) with an alpha of 0.025. Second, a 2×2 repeated-measures cluster permutation ANOVA [86] (for code see https://github.com/sashel/permANOVA) with the within-subject factors emotion (negative/neutral) and cognition (incongruent/congruent) was performed on t-values obtained from the first level, across subjects. To correct for multiple comparisons in four frequency bands (*θ*, *β*, *γ* and high *γ*) and two time windows (tROI Emo and tROI Flanker) we used Bonferroni correction with a factor of 8 leading to an alpha level of 0.00625. To correct for the number of grid voxels, we used a cluster-based correction approach [79] with 5000 permutations and a cluster-alpha level of 0.00625. Further, we obtained the coordinates of the active sources using the local maxima of the activation – without resorting to prior assumptions on their locations. This approach is in line with best practices recommendations for source reconstructions in MEG [87].

#### Temporal Generalization of condition classification

We used a time-resolved support vector machine analysis (t-SVM) with a linear kernel to test the ability of a classifier trained at each time sample to discriminate conditions at all other time samples [88] on source activity time courses as input data. The t-SVM was trained and tested separately for each subject, and each IFG subregion (IFG_Tri_, IFG_Op_ and IFG_Orb_). First, we z-normalized each source activity time course relative to the baseline period (see Temporal ROI definition for definition of baseline). Then, we smoothed the data with a Gaussian Kernel of ±25 ms and down sampled it to 250 Hz, similar to the approach in [89], and low-pass filtered at 60 Hz, in order to reduce the computational cost and increase the stimulus-to-noise ratio. After preprocessing, the source data were randomly assigned to one of twenty *supertrials* (per condition) and averaged (MATLAB code was adopted from [90], which builds on the library for SVMs by [91]). Finally, we separated these *supertrials* into training and testing data, with one *supertrial* per condition serving as test data and all others as training data. The binary classification was performed for each time point from 0 ms to 1000 ms after the onset of the emotional picture. Training one classifier at time t and generalizing it to time t’ was performed within the cross-validation so that t and t’ data came from independent sets of trials. To obtain a more robust estimate of the classification accuracy, we performed 200 iterations of *supertrials* creation, averaging and classification. The final temporal generalization classification reflects the average across these iterations. Classification for the emotional effects was performed between negative and neutral conditions, for cognitive effects classification was performed between incongruent and congruent and for the emotion-cognition interaction effects classification was performed between the pooled conditions of incongruent negative and congruent neutral versus the pooled conditions of incongruent neutral and congruent negative. This allowed to create two conditions with pooled data reflecting the interaction effect and thereby be classified with an SVM. For statistical testing, we employed a dependent-samples permutation t-test and a cluster-based correction method [79], where the number of permutations was set to 10000. The null hypothesis of no experimental effect for the two-dimensional classification matrix was equal to 50% chance level and tested within a tROI of 1000 ms including the emotional picture presentation and 500 ms of the Flanker stimulus presentation (tROI EMO and tROI Flanker) for the E contrast, while during the tROI Flanker for C and EC contrasts. The statistical tests were two-sided (the *α* value was set to 0.025) and Bonferroni corrected for the number of tests performed (nine tests; three IFG regions by three classifications: E, C and EC). Since the results of statistical analyses are random variables themselves, we additionally conducted bootstrapping of the cluster parameter estimates to obtain confidence intervals. Specifically, we calculated bootstrap-confidence intervals for maximum classification accuracy and the respective time by running the cluster statistics with 5000 bootstrap samples. We calculated point estimates (mean) and 95% confidence intervals from these bootstrap samples.

#### Non-parametric Granger Causality Analysis

To compute conditional Granger Causality (cGC), we employed a multivariate non-parametric spectral matrix factorization. We computed the CSD matrix of the source stimulus using a fast Fourier transform in combination with multitapers (5 Hz smoothing). We used the non-parametric variant of cGC [92] in order to avoid choosing a multivariate autoregressive model order, which can introduce a bias. Additionally, we used a block-wise approach [93] considering the first two principal components (PCs) of each source stimulus as a block, then estimating the cGC that a source X exerts over a source Y conditional on the remaining areas [30]. We applied the cGC to the three IFG subdivisions (IFG_Tri_, IFG_Orb_ and IFG_Op_) and the posterior sources that showed significant interaction effects: V2 and Precuneus (High *γ* band). We focused on bidirectional connectivity within IFG (IFG_Tri_, IFG_Op_ and IFG_Orb_) and of each IFG subdivision and V2 and Precuneus, respectively, in the post cognitive stimulus window (0.05 to 0.500 s). To alleviate the problem of non-stationarity we computed the cGC over two shorter time windows with 250 ms duration, with a spectral resolution of 2 Hz. The two time-sliding windows had an overlap of 50 ms: from 50 to 300 and from 250 to 500 ms. As the frequency ranges observed by [11] in relation to behavioral inhibition, were found to be low *γ* range, we broadened the statistical analysis from 10 to 44 Hz, despite our main interest lying on the *β*-band.

#### Statistical testing of cGC

First, we assessed whether the average cGC (in the frequency range of 8 − 44 Hz) of the source-target pairs was reliably above the bias level for each condition (negative congruent, neutral congruent, negative incongruent and neutral incongruent) separately. In order to estimate the bias, we randomly permuted the trials 200 times in each condition to create a surrogate distribution of mean cGC values. We tested if the found cGC value was in the upper 99.37% extreme (equivalent p*<*0.025 with Bonferroni correction for four conditions) of the surrogates’ distribution. If the average cGC exceeded the bias level, this source-target link was considered significant. These steps were repeated for each subject separately. Second, for both source-target pairs, we computed E, C and EC contrasts on cGC values at group level on subjects showing a significant link at least in one condition. The statistical comparison was performed in the range of 10 − 44 Hz using a dependent-samples permutation ANOVA. A cluster-based correction was used to account for multiple comparisons across frequencies [79]. Adjacent frequency samples with uncorrected p-values of 0.05 were considered as clusters. Fifty-thousand permutations were performed, and the critical *α* value was set to 0.05. Further Bonferroni correction was applied to account for multiple comparisons across links. Additionally, we report point estimates (mean) and 95% confidence intervals from bootstrapping with 50000 random resampling for each contrast: emotion (negative − neutral), cognition (incongruent − congruent) and emotion-cogniton interaction ((negative incongruent − neutral incongruent) − (negative congruent − neutral congruent)) for each link and each significant frequency range.

#### Correlation of top-down Granger Causality with behavior

To understand whether the links we found are of behavioral relevance, we used Bayesian linear regressions to determine which of the significant links from IFG to visual areas is the most informative factor to predict mean reaction time and mean accuracy, respectively. In the following, we describe the respective Bayesian model definition for predicting mean reaction time and mean accuracy data based on the links that showed significant effects of emotion, cognition or emotion-cognition interaction in Granger causality at certain frequency range: For the *j* th subject, we define the likelihood of the reaction time or the accuracy respectively given by y; denoted as:

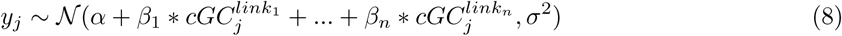

where for the *j* th subject, *α* is the intercept and encodes the grand average of reaction time or accuracy, respectively, and *β*_1_*…β_n_* capture the relation between the cGC links of the IFG subdivisions to Precuneus or V2 and the dependent variables (reaction time or accuracy). Additionally, we performed Bayesian regression regressing the differences in behavioral measures (Δ*^rt^* for reaction time, Δ*^acc^* for accuracy) on the difference in Granger causality (ΔcGC) across links selected based on the analysis above (links that had a 94% HDI of entirely one sign). Differences between conditions were computed as incongruent − congruent, in relation to cognitive processing (c contrast), and negative − neutral for emotional processing (E contrast). In the following, we describe the respective Bayesian model definition for a regression of Δ*^rt^* reaction time and Δ*^acc^* accuracy data:

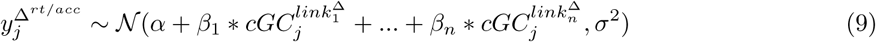

where for the Δ*^rt^* in the C contrast (incongruent − congruent) the selected *links* were the IFG_Orb_ to Precuneus and V2, while only the link IFG_Op_ to V2 entered the E contrast (negative − neutral) for Δ*^rt^* and Δ*^acc^*. We *z* -normalized all variables before modeling, and *β* coefficients are reported with respect to these standardized variables. All parameters were assumed to be drawn from normal distributions. We used non-informative priors in both models that can be found in.

### Software and settings for parameter inference of Bayesian models when using Markov chain Monte Carlo sampling

We estimated all the model regression coefficients using Bayesian inference with MCMC, using the python package PyMC3 [94] with NUTS (NO-U-Turn Sampling), using multiple independent Markov Chains. We ran four chains with 3000 burn-in (tuning) steps using NUTS. Then, each chain performed 8000 steps for the Bayesian Hierarchical model of reaction times and accuracy and 10000 steps for the Bayesian linear regression of *β* power and cGC values; those steps were used to approximate the posterior distribution. To check the validity of the sampling, we verified that the R-hat statistic was below 1.05. Finally, to assess the sensitivity of the posterior distribution of the estimated parameters to the choice of priors, we repeated the Bayesian analysis of behavioral and neural data using priors ten times broader (see [95]). The sensitivity analysis can be found in Supplementary Results of EEG source reconstruction). The Bayesian Hierarchical analysis of behavioral data and cGC values were conducted using Python (Version 3.7) [96] in Jupyter Notebook [97]. The code for fitting these models is freely available at OSF repository (https://osf.io/nrcuw/).

## Author contribution

Conceptualization, M.W. and O.T.; Methodology, A.D., E.P., Y.C.C., M.W.; Investigation, A.D., E.P., Y.C.C.; Writing – Original Draft, A.D.; Writing – Review Editing, A.D., E.P., Y.C.C., F.M.D., M.W., O.T.; Funding Acquisition, M.W. and O.T.; Resources, M.W. and O.T.; Supervision, M.W. and O.T.

## Acknowledgments

This project was funded by German Research Foundation (CRC 1193, project C04, MW and OT). MW, Volkswagen Stiftung, Award ID: Big Data in den Lebenswissenschaften. MW, Niedersächsisches Ministerium für Wissenschaft und Kultur (https://dx.doi.org/10.13039/501100010570), Award ID: Niedersächsisches Vorab. OT This work was supported by JST Moonshot RD Grant Number JPMJMS2292. We thank Shirin Hagner, Kristin Weineck, Sarah Brosche and Svenja Hummel for their help in data collection.

## Supporting information

### Supplementary information about behavioral effects of emotion-cognition interaction

The average reaction time was 547.75 ± 51.74*ms*. The increased cognitive load during incongruent versus congruent trials slowed down the reaction time by 95.2 ms on average. The increased emotional interference during negative compared to neutral trials slowed down the reaction time by 8.2 ms on average. The mean reaction time were 601.6 ± 59.49*ms* for negative incongruent trials, 593.6 ± 58.96*ms* for neutral incongruent trials, 506.6 ± 44.55*ms* for negative congruent trials and 498.2 ± 43.97*ms* for neutral congruent trials. There was a stronger emotional interference when the cognitive load was lower, as the difference on incongruent minus congruent for negative was 90.4 ms compared to 95.4 ms for neutral trials. This means subjects are slowed down in their reaction time due to higher emotional valence (negative) especially when the cognitive load is lower (congruent Flanker). For reaction time, a linear mixed effect model was used to predict reaction time based on an interaction of emotion (i.e., negative/neutral) and cognition (incongruent/congruent) as main effects, allowing for different intercepts per subject (random effects). For accuracy, the same model was used in a general linear mixed effect model approach. Both models were statistically tested against the respective null model, the same model but without the interaction term. To test for statistical significance of the interaction term, a maximum likelihood ratio approach [98] was used to compare the models. Model coding was done in R [99] with the lmer and glmer function from the lme4-package [100]. To test whether modeling the interaction is more informative compared to the null model without the interaction when predicting the reaction time, a maximum likelihood ratio was computed. The interaction of emotional load and cognitive load is significant (p=0.001563) in terms of reaction time (see). The accuracy was generally high, as the average accuracy was 95.18 ± 4.79% suggesting an easy feasibility of the task. Nevertheless, the increased cognitive load of incongruent compared to congruent trials led to a drop in accuracy of 7.15 % on average. The increased emotional interference during negative compared to neutral trials led to a drop in accuracy of 0.55 % on average. The mean accuracy was 91.2 ± 7.14% for negative incongruent, 92.0 ± 7.38% for neutral incongruent, 98.6 ± 2.53% for negative congruent and 98.9 ± 2.1% for neutral congruent trials. There was a stronger emotional interference when the cognitive load was higher, as the difference on incongruent minus congruent for negative was −7.4% compared to −6.8% for neutral. This means subjects make more incorrect responses during high cognitive load (incongruent Flanker) especially with high emotional load (negative valence). In sum, the cognitive effect was ten times stronger than the emotional effect, which in turn was ten times stronger than the interaction effect on the level of behavior (see Fig 1). To test whether modeling the interaction is more informative compared to the null model without the interaction when predicting the accuracy, a maximum likelihood ratio was computed. The interaction of emotional load and cognitive load is not significant (p=0.2199) in terms of accuracy (see).

**S1 Table.** Frequentist linear mixed models for reaction times and accuracy. Reaction time (RT) and accuracy (Acc) were predicted on a null model including only main effects and an interaction model including an interaction effect. Models were compared to test whether the interaction was informative.

| Model | AIC | BIC | deviance | $\chi^2$ | $\chi$ df | p |
| --- | --- | --- | --- | --- | --- | --- |
| RT Null Model | -350810 | -350761 | -350820 | NA | NA | NA |
| RT Interaction Model | -350818 | -350759 | -350830 | 10.002 | 1 | 0.001563** |
| Acc Null Model | 41713 | 41752 | 41705 | NA | NA | NA |
| Acc Interaction Model | 41714 | 41762 | 41704 | 1.5052 | 1 | 0.2199 |

**S2 Table.** Priors of Hierarchical Bayesian linear regression for reaction time and accuracy. Mean reaction time and mean accuracy per subject was predicted based on E, C and EC contrast (see Bayesian analysis of behavioral data)

| parameter | value |
| --- | --- |
| $\alpha_{intercept}$ | *Normal* ($\mu = 0, \sigma = 5$) |
| $\beta_E$ | *Normal*($\mu = 0, \sigma = 3$) |
| $\beta_C$ | *Normal*($\mu = 0, \sigma = 3$) |
| $\beta_{EC}$ | *Normal*($\mu = 0, \sigma = 3$) |
| $\alpha_{offset}$ | *Normal*($\mu = 0, \sigma = 1$) |
| $\beta_{E_{offset}}$ | *Normal*($\mu = 0, \sigma = 1$) |
| $\beta_{C_{offset}}$ | *Normal*($\mu = 0, \sigma = 1$) |
| $\sigma$ | *HalfNormal* ($\sigma = 1$) |

### Supplementary Results of EEG source reconstruction

#### 0.0.1 Definition of frequency bands of interest at the sensor level

To identify frequency bands of interest for the subsequent source-level analysis, we computed the contrast task against baseline at the sensor level, and identified four frequency bands of interest: *θ* 7 ± 2 Hz, *β* 21 ± 12 Hz, *γ* 49 ± 16 Hz and high *γ* 102 ± 38 Hz (center frequency ± spectral smoothing) for both time windows of interest before (tROI Emo) and after (tROI Flanker) ().

**S3 Figure.**
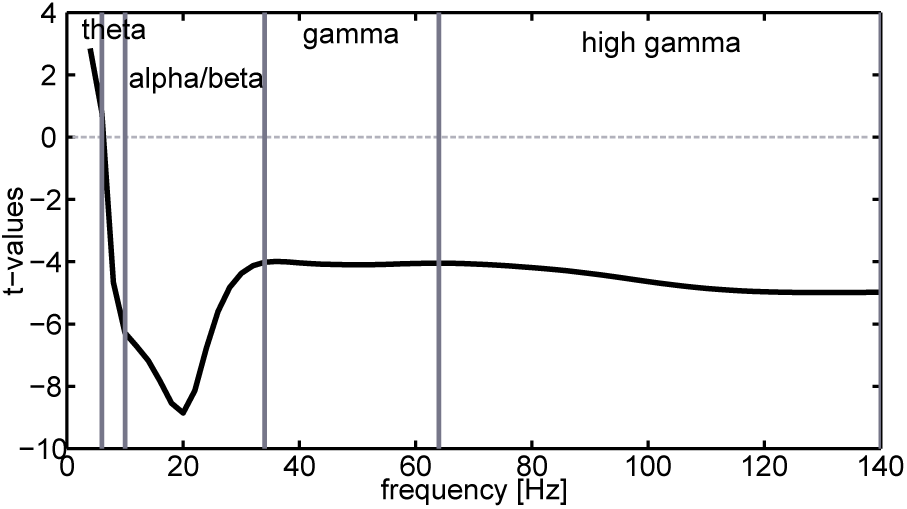
Spectral sensor level analysis for the task-independent definition of frequency bands of interest. Power spectrum analysis, obtained from pooled task conditions versus baseline to determine frequency bands (n=103). Grey lines in the power spectrum frame the frequency ranges of interest for subsequent beamforming analysis.

#### Supplementary results of source localization

In order to identify neural sources significantly modulated by the factors emotion and cognition in our task, and by their interaction, we reconstructed sources of each contrast (E, C and EC), both, before (tROI Emo) and after (tROI Flanker) Flanker signal onset. In we report the full results of four different frequency bands, *θ* 7 ± 2 Hz, *β* 21 ± 12 Hz, *γ* 49 ± 16 Hz and high *γ*102 ± 38 Hz (center frequency ± spectral smoothing) for both time windows of interest – before (tROI Emo) and after (tROI Flanker).

**S4 Figure.**
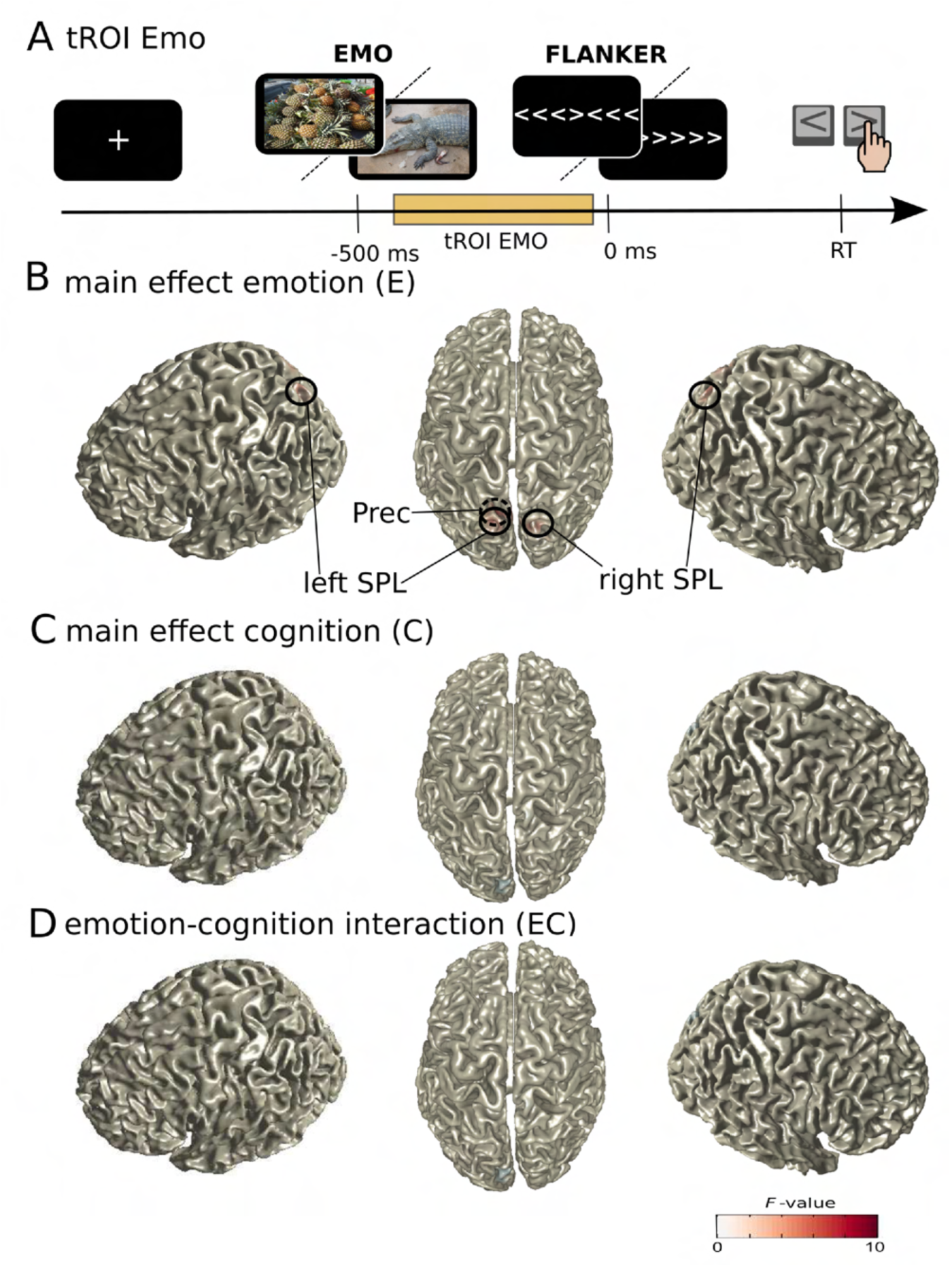
Source activity in the *θ* band (5-9 Hz) during tROI Emo. Surface plots show significant F-values of ANOVA (cluster-based permutation test, Bonferroni corrected for four frequency bands and two time windows, alpha and clusteralpha *<*0.00625, n=103). Peak voxels (local extrema) of significant clusters are indicated with circles and labels (for MNI coordinates, see). Dashed circles mark peak voxels invisible in this representation. A, Task design and analyzed tROI; B, main effect of emotion (E); C, main effect of cognition (C); D, interaction effect between emotion and cognition (EC).

**S5 Figure.**
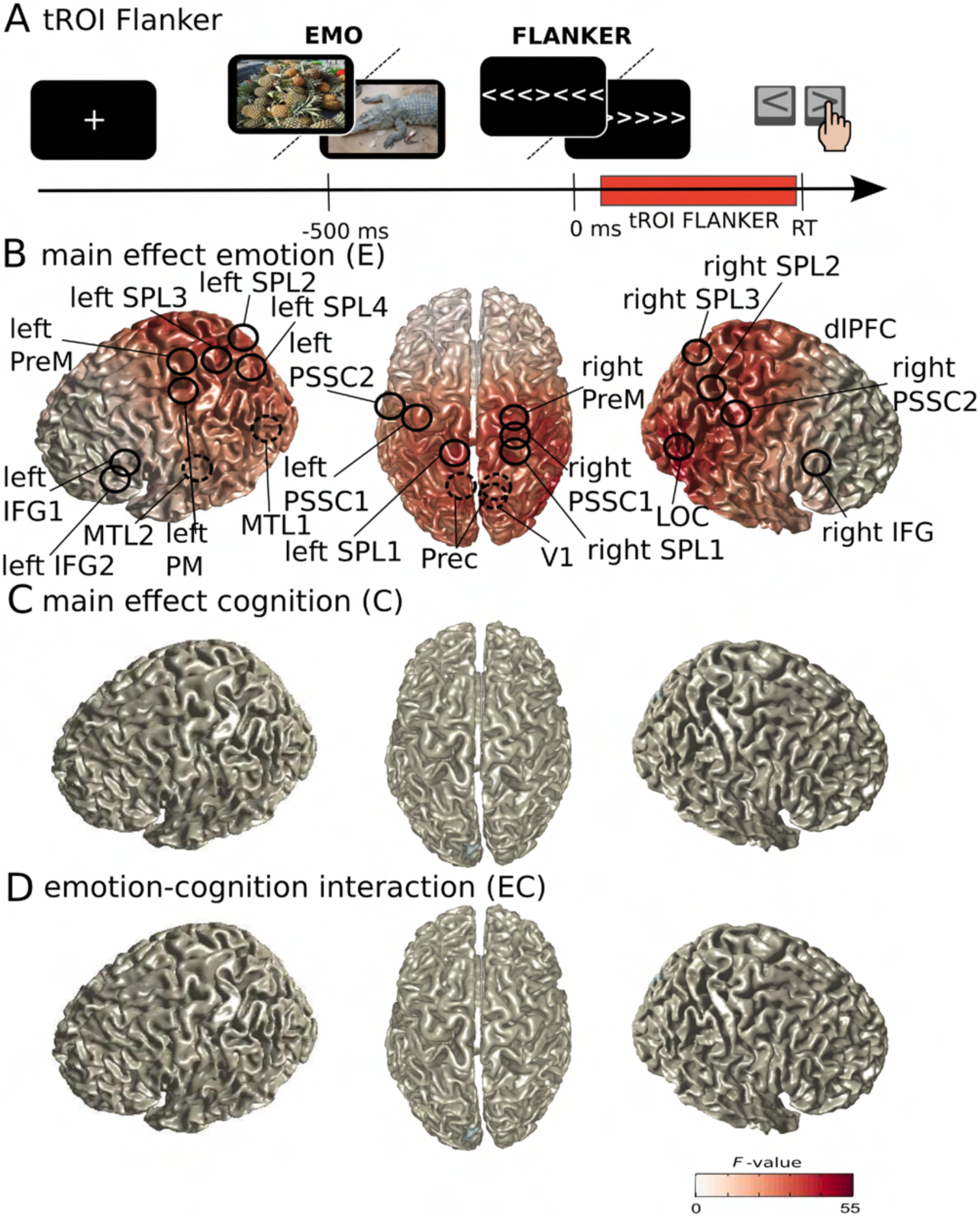
Source activity in the *θ* band (5-9 Hz) during tROI Flanker. Surface plots show significant F-values of ANOVA (cluster-based permutation test, Bonferroni corrected for four frequency bands and two time windows, alpha and clusteralpha *<*0.00625, n=103). Peak voxels (local extrema) of significant clusters are indicated with circles and labels (for MNI coordinates, see). Dashed circles mark peak voxels invisible in this representation. A, Task design and analyzed tROI; B, main effect of emotion (E); C, main effect of cognition (C); D, interaction effect between emotion and cognition (EC).

**S6 Figure.**
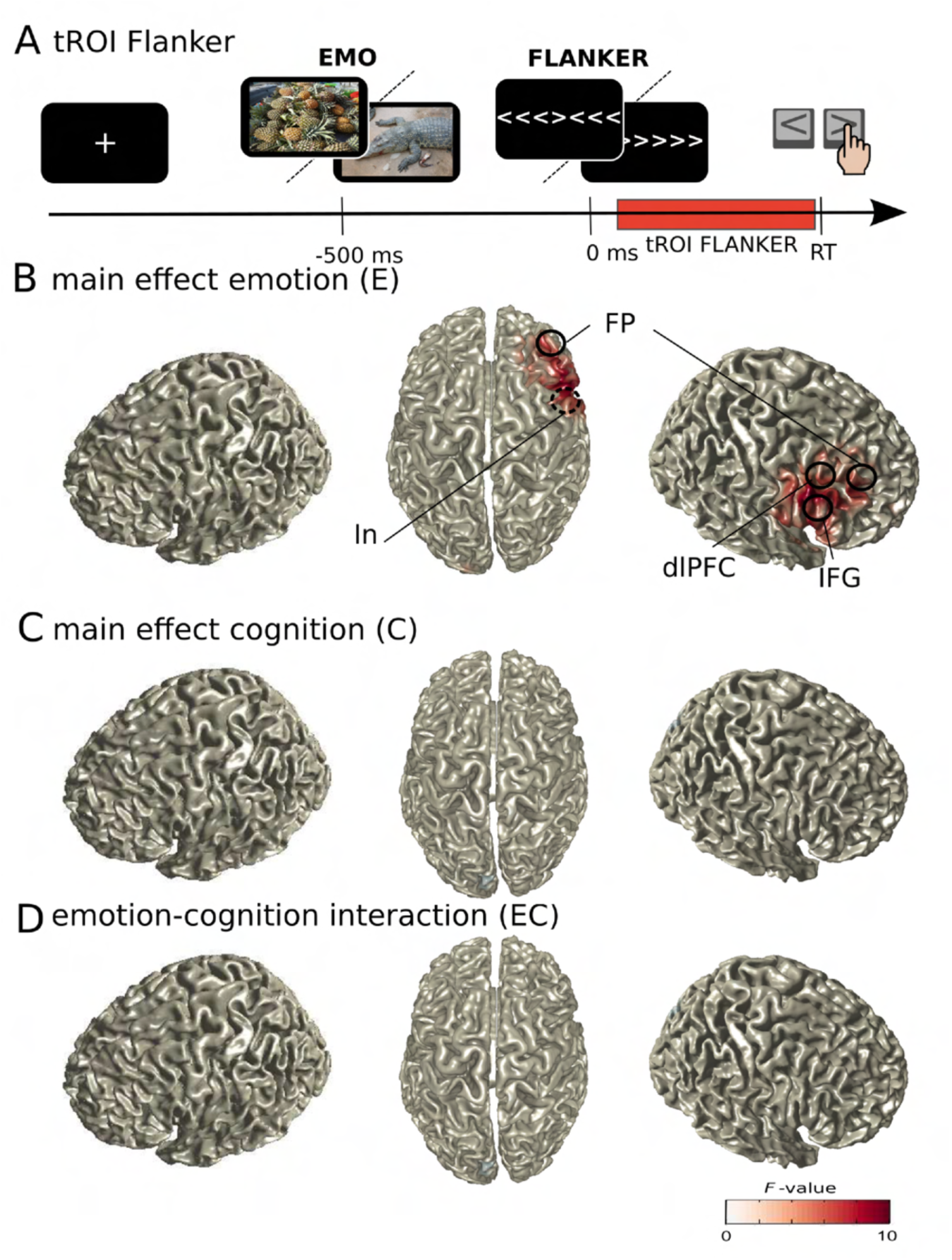
Source activity in the *γ* band (33-65 Hz) during tROI Flanker. Surface plots show significant F-values of ANOVA (cluster-based permutation test, Bonferroni corrected for four frequency bands and two time windows, alpha and clusteralpha *<*0.00625, n=103). Peak voxels (local extrema) of significant clusters are indicated with circles and labels (for MNI coordinates, see). Dashed circles mark peak voxels invisible in this representation. A, Task design and analyzed tROI; B, main effect of emotion (E); C, main effect of cognition (C); D, interaction effect between emotion and cognition (EC).

**S7 Figure.**
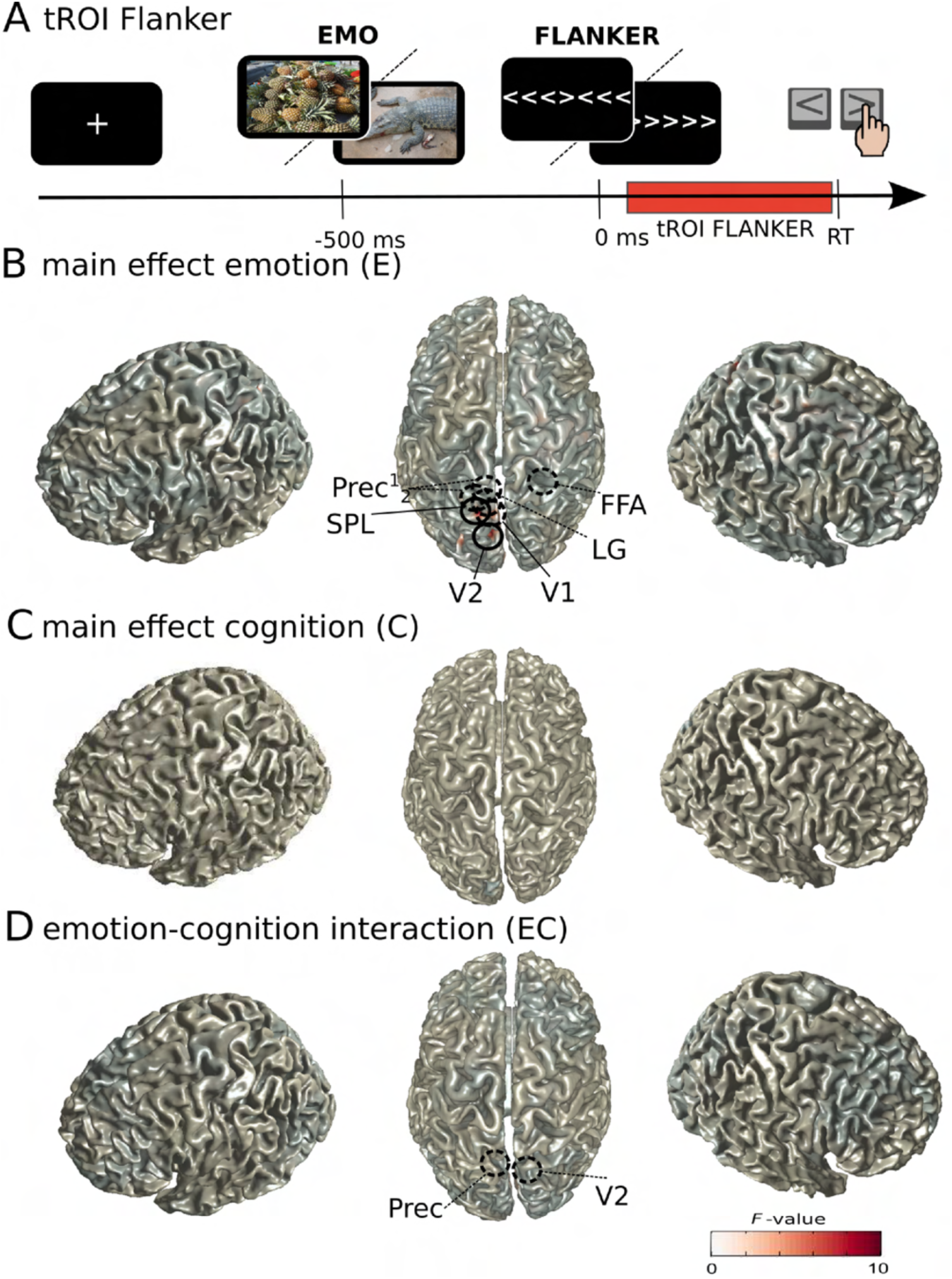
Source activity in the High *γ* band (64-140 Hz) during tROI Flanker. Surface plots show significant F-values of ANOVA (cluster-based permutation test, Bonferroni corrected for four frequency bands and two time windows, alpha and clusteralpha *<*0.00625, n=103). Peak voxels (local extrema) of significant clusters are indicated with circles and labels (for MNI coordinates, see). Dashed circles mark peak voxels invisible in this representation. A, Task design and analyzed tROI; B, main effect of emotion (E); C, main effect of cognition (C); D, interaction effect between emotion and cognition (EC).

**S8 Table.** Overview of significant sources. Significant sources for all frequency bands and tROIs are given for the contrasts (Contr.) of emotion (E), cognition (C) and emotion-cognition interaction (EC). The table shows sources (Region (label)) with respective coordinates and F-values. Furthermore, it shows the point estimate (p.e.) of source activity (mean difference for main effects mean double difference for interaction effects) and the bootstrap Confidence-Interval of the effect (95% CI). Additional Cluster F- and P-values are presented.

| Freq. | tROI. | Contr. | Region (label) | Coordinat.<br>(x,y,z) | Source<br>F-value | p.e.<br>(Mean) | Bootstrap<br>95% CI | Cluster<br>F-value | Cluster<br>P-value |
| --- | --- | --- | --- | --- | --- | --- | --- | --- | --- |
| $\theta$ | EMO | E | Superior Parietal Lobe (rSPL) R | 15 -80 60 | 11.1455 | -0.4158 | [-0.6641; -0.1692] | 11.1455 | 3.9992e-04 |
| $\theta$ | EMO | E | Precuneus (IPrec) L | -5 -70 40 | 8.9030 | -0.3417 | [-0.5697; -0.1146] | 33.0149 | 1.9996e-04 |
| $\theta$ | EMO | E | Superior Parietal Lobe (ISPL) L | -25 -90 50 | 8.3064 | -0.3503 | [-0.5858; -0.1043] | 8.3064 | 3.9992e-04 |
| $\theta$ | Flanker | E | Lateral Occipital Cortex (rLOC) R | 55 -70 10 | 46.6805 | -0.8177 | [-1.0558; -0.5825] | 6.1468e+04 | 1.9996e-04 |
| $\theta$ | Flanker | E | Superior Parietal Lobe (rSPL) R | 25 -50 40 | 44.4014 | -0.7868 | [-1.0194; -0.5505] | 6.1468e+04 | 1.9996e-04 |
| $\theta$ | Flanker | E | Primary Visual Cortex (rV1) R | 5 -80 10 | 43.9917 | -0.7519 | [-0.9778; -0.5291] | 6.1468e+04 | 1.9996e-04 |
| $\theta$ | Flanker | E | Primary Somatosensory Cortex R (rPSSC), BA2 | 25 -40 60 | 43.9502 | -0.7575 | [-0.9854; -0.5344] | 6.1468e+04 | 1.9996e-04 |
| $\theta$ | Flanker | E | Superior Parietal Lobe L (ISPL) | -15 -40 80 | 41.8829 | -0.7834 | [-1.0158; -0.5373] | 6.1468e+04 | 1.9996e-04 |
| $\theta$ | Flanker | E | Precuneous Cortex R (rPrec) | 15 -70 30 | 41.7265 | -0.7527 | [-0.9829; -0.5226] | 6.1468e+04 | 1.9996e-04 |
| $\theta$ | Flanker | E | Primary Somatosensory Cortex R (rPSSC), BA1 | 55 -20 60 | 41.3273 | -0.6751 | [-0.8769; -0.4609] | 6.1468e+04 | 1.9996e-04 |
| $\theta$ | Flanker | E | Premotor Cortex R (rPM) | 25 -20 80 | 41.1597 | -0.6969 | [-0.9070; -0.4762] | 6.1468e+04 | 1.9996e-04 |
| $\theta$ | Flanker | E | Precuneous Cortex L (IPrec) | -5 -70 30 | 41.0866 | -0.7976 | [-1.0435; -0.5524] | 6.1468e+04 | 1.9996e-04 |
| $\theta$ | Flanker | E | Superior Parietal Lobe L (ISPL) | -5 -80 60 | 40.6483 | -0.7424 | [-0.9750; -0.5121] | 6.1468e+04 | 1.9996e-04 |
| $\theta$ | Flanker | E | Inferior Parietal Lobe R (rIPL) | 55 -40 40 | 40.2498 | -0.7208 | [-0.9448; -0.4948] | 6.1468e+04 | 1.9996e-04 |
| $\theta$ | Flanker | E | Superior Parietal Lobe (rSPL) R | 15 -80 60 | 40.0227 | -0.7825 | [-1.0306; -0.5403] | 6.1468e+04 | 1.9996e-04 |
| $\theta$ | Flanker | E | Superior Parietal Lobe L (ISPL) | -15 -50 60 | 38.5924 | -0.7574 | [-0.9965; -0.5163] | 6.1468e+04 | 1.9996e-04 |
| $\theta$ | Flanker | E | Superior Parietal Lobe L (ISPL) | -25 -80 60 | 38.1349 | -0.7820 | [-1.0372; -0.5379] | 6.1468e+04 | 1.9996e-04 |
| $\theta$ | Flanker | E | Cerebellum R (rCer) | 15 -50 -30 | 37.2761 | -0.7205 | [-0.9513; -0.4818] | 6.1468e+04 | 1.9996e-04 |
| $\theta$ | Flanker | E | Cerebellum R (rCer) | 5 -80 -30 | 35.5559 | -0.7370 | [-0.9811; -0.4893] | 6.1468e+04 | 1.9996e-04 |
| $\theta$ | Flanker | E | Primary Somatosensory Cortex L (IPSSC) | -55 -20 60 | 34.9523 | -0.6195 | [-0.8256; -0.4114] | 6.1468e+04 | 1.9996e-04 |
| $\theta$ | Flanker | E | Primary Motor Cortex L (IPrM) | -35 -10 70 | 33.6669 | -0.6428 | [-0.8573; -0.4141] | 6.1468e+04 | 1.9996e-04 |
| $\theta$ | Flanker | E | Premotor Cortex L (IPM) | -15 -10 80 | 33.4476 | -0.6700 | [-0.8955; -0.4347] | 6.1468e+04 | 1.9996e-04 |
| $\theta$ | Flanker | E | Primary Somatosensory Cortex L (IPSSC) | -65 -10 40 | 28.6332 | -0.5728 | [-0.7849; -0.3628] | 6.1468e+04 | 1.9996e-04 |
| $\theta$ | Flanker | E | Middle Temporal Lobe L (ITL) | -65 -60 10 | 26.4450 | -0.6204 | [-0.8632; -0.3853] | 6.1468e+04 | 1.9996e-04 |
| $\theta$ | Flanker | E | Middle Temporal Lobe L (ITL) | -65 0 -10 | 18.5366 | -0.4800 | [-0.7096; -0.2655] | 6.1468e+04 | 1.9996e-04 |
| $\theta$ | Flanker | E | Inferior Frontal Gyrus R (rIFG) | 55 40 0 | 15.3509 | -0.4218 | [-0.6327; -0.2090] | 6.1468e+04 | 1.9996e-04 |
| $\theta$ | Flanker | E | Inferior Frontal Gyrus L (IFG) | -55 40 10 | 12.1197 | -0.3917 | [-0.6177; -0.1716] | 6.1468e+04 | 1.9996e-04 |
| $\theta$ | Flanker | E | Inferior Frontal Gyrus L (IFG) | -45 50 20 | 11.2494 | -0.3972 | [-0.6322; -0.1639] | 11.2494 | 1.9996e-04 |
| $\beta$ | Flanker | E | Fusiform Face Area R (rFFA) | 35 -50 10 | 16.1438 | 0.4319 | [0.2187; 0.6455] | 5.2295e+03 | <1.9996e-04 |
| $\beta$ | Flanker | E | dorsomedial Prefrontal Cortex R (rdmPFC) | 5 40 40 | 15.9367 | 0.4302 | [0.2212; 0.6483] | 5.2295e+03 | <1.9996e-04 |
| $\beta$ | Flanker | E | dorsolateral Prefrontal Cortex R (rdlPFC) | 25 30 60 | 13.5796 | 0.3963 | [0.1808; 0.6073] | 5.2295e+03 | <1.9996e-04 |
| $\beta$ | Flanker | E | lateral Occipital Cortex R (rLOC) | 35 -70 30 | 13.3564 | 0.3686 | [0.1680; 0.5689] | 5.2295e+03 | <1.9996e-04 |
| $\beta$ | Flanker | E | Frontal Pole R (rFP) | 5 60 40 | 12.8408 | 0.4005 | [0.1744; 0.6177] | 5.2295e+03 | <1.9996e-04 |
| $\beta$ | Flanker | E | Angular Gyrus R (rAG) | 45 -50 20 | 12.6571 | 0.3737 | [0.1714; 0.5859] | 5.2295e+03 | <1.9996e-04 |
| $\beta$ | Flanker | E | Superior Frontal Gyrus R (rSFG) | 25 10 60 | 12.4352 | 0.3686 | [0.1613; 0.5781] | 5.2295e+03 | <1.9996e-04 |
| $\beta$ | Flanker | E | Premotor Cortex R (rPM) | 45 0 50 | 11.2470 | 0.3553 | [0.1453; 0.5647] | 19.5099 | <1.9996e-04 |
| $\beta$ | Flanker | E | Inferior Frontal Gyrus R (rIFG) | 45 40 0 | 10.4409 | 0.3852 | [0.1510; 0.6221] | 5.2295e+03 | <1.9996e-04 |
| $\beta$ | Flanker | E | Precuneous Cortex R (rPrec) | 15 -70 50 | 9.7742 | 0.3459 | [0.1311; 0.5695] | 5.2295e+03 | <1.9996e-04 |
| $\beta$ | Flanker | E | Frontal Pole L (IFP) | -25 50 10 | 9.6548 | 0.3641 | [0.1238; 0.5947] | 25.9950 | <1.9996e-04 |
| $\beta$ | Flanker | E | Primary Somatosensory Cortex R (rPSSC) | 45 -10 30 | 8.8410 | 0.3268 | [0.1172; 0.5519] | 5.2295e+03 | <1.9996e-04 |
| $\beta$ | Flanker | E | Inferior Frontal Gyrus L (IFG) | -35 30 10 | 8.7014 | 0.3131 | [0.1040; 0.5265] | 8.7014 | <1.9996e-04 |
| $\beta$ | Flanker | E | Primary Motor Cortex R (rPrMC) | 35 -20 50 | 8.2433 | 0.3131 | [0.1016; 0.5327] | 8.2433 | <1.9996e-04 |
| $\beta$ | Flanker | C | dorsolateral Prefrontal Cortex R (rdlPFC) | 45 20 30 | 8.5053 | 0.3048 | [0.1082; 0.5220] | 8.5053 | <1.9996e-04 |
| $\beta$ | Flanker | C | medial Temporal Lobe R (rmTL) | 65 -10 -10 | 8.3118 | 0.3278 | [0.1194; 0.5716] | 8.3118 | <1.9996e-04 |
| $\beta$ | Flanker | C | Inferior Frontal Gyrus R (rIFG) | 55 10 10 | 7.9618 | 0.3034 | [0.1010; 0.5278] | 7.9618 | <1.9996e-04 |
| $\beta$ | Flanker | EC | Inferior Frontal Gyrus R (rIFG) | 45 40 20 | 9.6267 | -0.5948 | [-0.9589; -0.1966] | 9.6267 | <1.9996e-04 |
| $\gamma$ | Flanker | E | dorsolateral Prefrontal Cortex R (dlPFC) | 55 20 40 | 14.7569 | 0.3379 | [0.1642; 0.5134] | 586.3702 | <1.9996e-04 |
| $\gamma$ | Flanker | E | Inferior Frontal Gyrus R (IFG) | 65 20 10 | 14.1434 | 0.3717 | [0.1730; 0.5644] | 586.3702 | <1.9996e-04 |
| $\gamma$ | Flanker | E | orbitofrontal Frontal Pole R (oFP) | 35 50 30 | 12.9234 | 0.2990 | [0.1364; 0.4642] | 586.3702 | <1.9996e-04 |
| $\gamma$ | Flanker | E | Insula R (In) | 45 20 -10 | 12.7424 | 0.3383 | [0.1470; 0.5234] | 586.3702 | <1.9996e-04 |
| $\gamma$ | Flanker | E | ventromedial Frontal Pole R (vFP) | 15 60 0 | 9.2366 | 0.2885 | [0.1052; 0.4790] | 9.2366 | 0.0028 |
| High $\gamma$ | Flanker | E | Superior Parietal Lobe L (ISPL) | -15 -70 60 | 10.5863 | -0.2977 | [-0.4814; -0.1177] | 10.5863 | <1.9996e-04 |
| High $\gamma$ | Flanker | E | Secondary Visual Cortex L (IV2) | -5 -80 30 | 9.6571 | -0.2956 | [-0.4866; -0.1086] | 17.8269 | <1.9996e-04 |
| High $\gamma$ | Flanker | E | Primary Visual Cortex V1 L (IV1) | -5 -80 10 | 8.9396 | -0.3073 | [-0.5139; -0.1080] | 8.9396 | <1.9996e-04 |
| High $\gamma$ | Flanker | E | Lingual Gyrus L (ILG) | -5 -70 -10 | 8.4952 | -0.2811 | [-0.4740; -0.0921] | 8.4952 | <1.9996e-04 |
| High $\gamma$ | Flanker | E | Fusiform Face Area R (rFFA) | 25 -50 10 | 8.4180 | -0.2715 | [-0.4545; -0.0857] | 8.4180 | <1.9996e-04 |
| High $\gamma$ | Flanker | E | Precuneous Cortex L 1 (IPrec) | -5 -60 10 | 8.3565 | -0.2789 | [-0.4714; -0.0882] | 8.3565 | <1.9996e-04 |
| High $\gamma$ | Flanker | E | Precuneous Cortex L 2 (IPrec) | -5 -60 30 | 8.3055 | -0.2913 | [-0.4961; -0.0943] | 8.3055 | <1.9996e-04 |
| High $\gamma$ | Flanker | EC | Secondary Visual Cortex R (rV2) | 5 -80 20 | 7.9948 | -0.4994 | [-0.8461; -0.1485] | 7.9948 | <1.9996e-04 |
| High $\gamma$ | Flanker | EC | Precuneous Cortex L (IPrec) | -5 -60 30 | 7.8551 | -0.4667 | [-0.7983; -0.1334] | 7.8551 | <1.9996e-04 |

#### 0.0.2 Supplementary information on Bayesian priors

**S9 Table.** Priors of Bayesian linear regression between reaction time, accuracy and cGC. Mean reaction time and mean accuracy per subject were predicted based on cGC of each significant link of respective significant frequencies in the respective significant E, C, EC contrast (see Correlation of top-down Granger Causality with behavior and Fig7).

| parameter | value |
| --- | --- |
| $\alpha_{intercept}$ | *Normal* ($\mu = 0, \sigma = 0.5$) |
| $\beta_{links}$ | *Normal*($\mu = 0, \sigma = 0.5$) |
| $\beta_{links\Delta}$ | *Normal*($\mu = 0, \sigma = 0.5$) |
| $\sigma$ | *HalfNormal* ($\sigma = 1$) |

#### 0.0.3 Supplementary information on spectrally-resolved cGC from IFG to Precuneus and V2

**S10 Figure.**
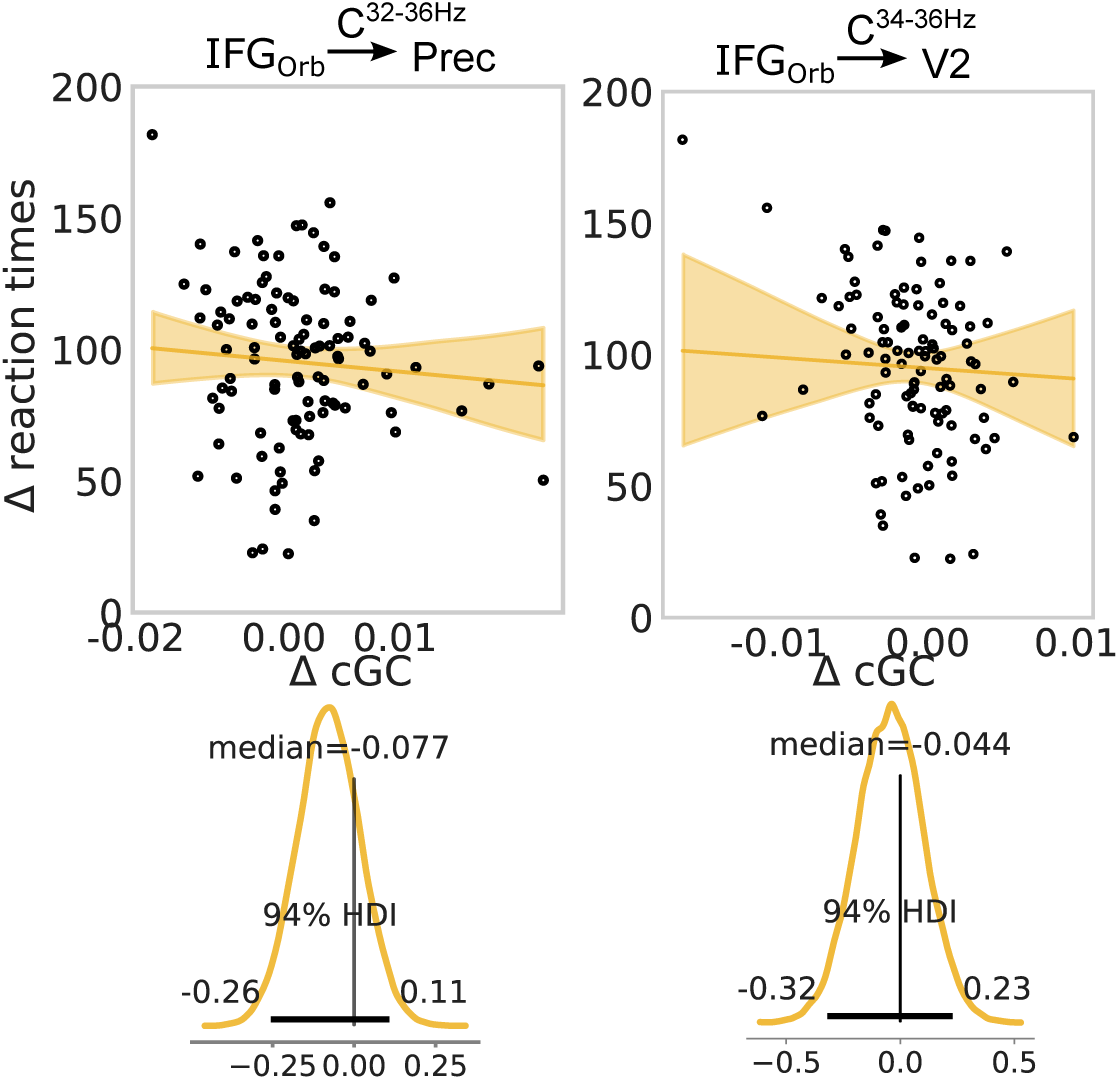
Spectrally-resolved cGC from IFG to Precuneus and V2 correlates with behavior. Linear regression for a regression of the differences between conditions (C) in behavior (reaction times) on the differences between conditions for cGC (ΔcGC), for cGC from IFG_Orb_ to Prec and V2. Corresponding marginal posterior distributions are shown below. See Correlation of top-down Granger Causality with behavior for a detailed description

## References

1. Okon-Singer H, Hendler T, Pessoa L, Shackman AJ. The neurobiology of emotion–cognition interactions: fundamental questions and strategies for future research. Frontiers in human neuroscience. 2015;9:58.

2. Carretié L. Exogenous (automatic) attention to emotional stimuli: a review. Cognitive, Affective, & Behavioral Neuroscience. 2014;14:1228–1258.

3. Pessoa L. How do emotion and motivation direct executive control? Trends in cognitive sciences. 2009;13(4):160–166.

4. Pessoa L. Précis on the cognitive-emotional brain. Behavioral and Brain Sciences. 2015;38.

5. Miyake A, Friedman NP, Emerson MJ, Witzki AH, Howerter A, Wager TD. The unity and diversity of executive functions and their contributions to complex “frontal lobe” tasks: A latent variable analysis. Cognitive psychology. 2000;41(1):49–100.

6. Friedman NP, Miyake A. The relations among inhibition and interference control functions: a latent-variable analysis. Journal of experimental psychology: General. 2004;133(1):101.

7. Etkin A, Schatzberg AF. Common abnormalities and disorder-specific compensation during implicit regulation of emotional processing in generalized anxiety and major depressive disorders. American Journal of Psychiatry. 2011;168(9):968–978.

8. Kronhaus DM, Lawrence NS, Williams AM, Frangou S, Brammer MJ, Williams SC, et al. Stroop performance in bipolar disorder: further evidence for abnormalities in the ventral prefrontal cortex. Bipolar disorders. 2006;8(1):28–39.

9. Bremner JD, Vermetten E, Vythilingam M, Afzal N, Schmahl C, Elzinga B, et al. Neural correlates of the classic color and emotional stroop in women with abuse-related posttraumatic stress disorder. Biological psychiatry. 2004;55(6):612–620.

10. Vorwerk J, Oostenveld R, Piastra MC, Magyari L, Wolters CH. The FieldTrip-SimBio pipeline for EEG forward solutions. Biomedical engineering online. 2018;17(1):1–17.

11. Schaum M, Pinzuti E, Sebastian A, Lieb K, Fries P, Mobascher A, et al. Right inferior frontal gyrus implements motor inhibitory control via beta-band oscillations in humans. Elife. 2021;10:e61679.

12. Sebastian A, Jung P, Neuhoff J, Wibral M, Fox PT, Lieb K, et al. Dissociable attentional and inhibitory networks of dorsal and ventral areas of the right inferior frontal cortex: a combined task-specific and coordinate-based meta-analytic fMRI study. Brain Structure and Function. 2016;221:1635–1651.

13. Stahl C, Voss A, Schmitz F, Nuszbaum M, Tüscher O, Lieb K, et al. Behavioral components of impulsivity. Journal of Experimental Psychology: General. 2014;143(2):850.

14. Larson J, Mavrogordatos T. The Jaynes–Cummings Model and Its Descendants: Modern research directions. IoP Publishing; 2021.

15. Hartwigsen G, Neef NE, Camilleri JA, Margulies DS, Eickhoff SB. Functional segregation of the right inferior frontal gyrus: evidence from coactivation-based parcellation. Cerebral Cortex. 2019;29(4):1532–1546.

16. Boen R, Raud L, Huster RJ. Inhibitory Control and the Structural Parcelation of the Right Inferior Frontal Gyrus. Frontiers in Human Neuroscience. 2022;16.

17. Wang C, Rajagovindan R, Han SM, Ding M. Top-down control of visual alpha oscillations: sources of control signals and their mechanisms of action. Frontiers in human neuroscience. 2016;10:15.

18. Pessoa L, Padmala S, Kenzer A, Bauer A. Interactions between cognition and emotion during response inhibition. Emotion. 2012;12(1):192.

19. Sebastian A, Pohl M, Klöppel S, Feige B, Lange T, Stahl C, et al. Disentangling common and specific neural subprocesses of response inhibition. Neuroimage. 2013;64:601–615.

20. Hannah R, Aron AR. Towards real-world generalizability of a circuit for action-stopping. Nature Reviews Neuroscience. 2021;22(9):538–552.

21. Underwood R, Tolmeijer E, Wibroe J, Peters E, Mason L. Networks underpinning emotion: A systematic review and synthesis of functional and effective connectivity. NeuroImage. 2021;243:118486.

22. Langner R, Leiberg S, Hoffstaedter F, Eickhoff SB. Towards a human self-regulation system: Common and distinct neural signatures of emotional and behavioural control. Neuroscience & Biobehavioral Reviews. 2018;90:400–410.

23. Modha DS, Singh R. Network architecture of the long-distance pathways in the macaque brain. Proceedings of the National Academy of Sciences. 2010;107(30):13485–13490.

24. Paquola C, Amunts K, Evans A, Smallwood J, Bernhardt B. Closing the mechanistic gap: the value of microarchitecture in understanding cognitive networks. Trends in cognitive sciences. 2022;26(10):873–886.

25. Song S, Zilverstand A, Song H, d’Oleire Uquillas F, Wang Y, Xie C, et al. The influence of emotional interference on cognitive control: A meta-analysis of neuroimaging studies using the emotional Stroop task. Scientific reports. 2017;7(1):1–9.

26. Verbruggen F, Aron AR, Stevens MA, Chambers CD. Theta burst stimulation dissociates attention and action updating in human inferior frontal cortex. Proceedings of the National Academy of Sciences. 2010;107(31):13966–13971.

27. Tops M, Boksem MA. A potential role of the inferior frontal gyrus and anterior insula in cognitive control, brain rhythms, and event-related potentials. Frontiers in psychology. 2011;2:330.

28. Corbetta M, Patel G, Shulman GL. The reorienting system of the human brain: from environment to theory of mind. Neuron. 2008;58(3):306–324.

29. Bibbig A, Traub RD, Whittington MA. Long-range synchronization of *γ* and *β* oscillations and the plasticity of excitatory and inhibitory synapses: a network model. Journal of Neurophysiology. 2002;88(4):1634–1654.

30. Bastos AM, Vezoli J, Bosman CA, Schoffelen JM, Oostenveld R, Dowdall JR, et al. Visual areas exert feedforward and feedback influences through distinct frequency channels. Neuron. 2015;85(2):390–401.

31. Brovelli A, Ding M, Ledberg A, Chen Y, Nakamura R, Bressler SL. Beta oscillations in a large-scale sensorimotor cortical network: directional influences revealed by Granger causality. Proceedings of the National Academy of Sciences. 2004;101(26):9849–9854.

32. Popov T, Westner BU, Silton RL, Sass SM, Spielberg JM, Rockstroh B, et al. Time course of brain network reconfiguration supporting inhibitory control. Journal of Neuroscience. 2018;38(18):4348–4356.

33. Yang K, Tong L, Shu J, Zhuang N, Yan B, Zeng Y. High gamma band EEG closely related to emotion: evidence from functional network. Frontiers in human neuroscience. 2020;14:89.

34. Benchenane K, Tiesinga PH, Battaglia FP. Oscillations in the prefrontal cortex: a gateway to memory and attention. Current opinion in neurobiology. 2011;21(3):475–485.

35. Norman DA, Bobrow DG. On data-limited and resource-limited processes. Cognitive psychology. 1975;7(1):44–64.

36. Desimone R, Duncan J, et al. Neural mechanisms of selective visual attention. Annual review of neuroscience. 1995;18(1):193–222.

37. Lavie N, Hirst A, De Fockert JW, Viding E. Load theory of selective attention and cognitive control. Journal of experimental psychology: General. 2004;133(3):339.

38. Botvinick MM, Cohen JD, Carter CS. Conflict monitoring and anterior cingulate cortex: an update. Trends in cognitive sciences. 2004;8(12):539–546.

39. Carter CS, Braver TS, Barch DM, Botvinick MM, Noll D, Cohen JD. Anterior cingulate cortex, error detection, and the online monitoring of performance. Science. 1998;280(5364):747–749.

40. Bush G, Luu P, Posner MI. Cognitive and emotional influences in anterior cingulate cortex. Trends in cognitive sciences. 2000;4(6):215–222.

41. Allman JM, Hakeem A, Erwin JM, Nimchinsky E, Hof P. The anterior cingulate cortex: the evolution of an interface between emotion and cognition. Annals of the New York Academy of Sciences. 2001;935(1):107–117.

42. Nigg JT. Attention-deficit/hyperactivity disorder: Endophenotypes, structure, and etiological pathways. Current Directions in Psychological Science. 2010;19(1):24–29.

43. Christ SE, Kester LE, Bodner KE, Miles JH. Evidence for selective inhibitory impairment in individuals with autism spectrum disorder. Neuropsychology. 2011;25(6):690.

44. Nigg JT, Silk KR, Stavro G, Miller T. Disinhibition and borderline personality disorder. Development and psychopathology. 2005;17(4):1129–1149.

45. Carver CS, Johnson SL, Joormann J. Serotonergic function, two-mode models of self-regulation, and vulnerability to depression: what depression has in common with impulsive aggression. Psychological bulletin. 2008;134(6):912.

46. Fineberg NA, Potenza MN, Chamberlain SR, Berlin HA, Menzies L, Bechara A, et al. Probing compulsive and impulsive behaviors, from animal models to endophenotypes: a narrative review. Neuropsychopharmacology. 2010;35(3):591–604.

47. Groman SM, James AS, Jentsch JD. Poor response inhibition: at the nexus between substance abuse and attention deficit/hyperactivity disorder. Neuroscience & Biobehavioral Reviews. 2009;33(5):690–698.

48. Serra-Blasco M, Radua J, Soriano-Mas C, Gómez-Benlloch A, Porta-Casteràs D, Carulla-Roig M, et al. Structural brain correlates in major depression, anxiety disorders and post-traumatic stress disorder: A voxel-based morphometry meta-analysis. Neuroscience & Biobehavioral Reviews. 2021;129:269–281.

49. Hollunder B, Ostrem JL, Sahin IA, Rajamani N, Oxenford S, Butenko K, et al. Segregating the Frontal Cortex with Deep Brain Stimulation. medRxiv. 2023; p. 2023–03.

50. Puiu AA, Wudarczyk O, Kohls G, Bzdok D, Herpertz-Dahlmann B, Konrad K. Meta-analytic evidence for a joint neural mechanism underlying response inhibition and state anger. Human brain mapping. 2020;41(11):3147–3160.

51. Zhu J, Li J, Li X, Rao J, Hao Y, Ding Z, et al. Neural basis of the emotional conflict processing in major depression: ERPs and source localization analysis on the N450 and P300 components. Frontiers in human neuroscience. 2018;12:214.

52. Zrenner B, Zrenner C, Gordon PC, Belardinelli P, McDermott EJ, Soekadar SR, et al. Brain oscillation-synchronized stimulation of the left dorsolateral prefrontal cortex in depression using real-time EEG-triggered TMS. Brain stimulation. 2020;13(1):197–205.

53. Zrenner C, Belardinelli P, Müller-Dahlhaus F, Ziemann U. Closed-loop neuroscience and non-invasive brain stimulation: a tale of two loops. Frontiers in cellular neuroscience. 2016;10:92.

54. Schunk D, Berger EM, Hermes H, Winkel K, Fehr E. Teaching self-regulation. Nature Human Behaviour. 2022;6(12):1680–1690.

55. Sheehan DV, Lecrubier Y, Sheehan KH, Amorim P, Janavs J, Weiller E, et al. The Mini-International Neuropsychiatric Interview (MINI): the development and validation of a structured diagnostic psychiatric interview for DSM-IV and ICD-10. Journal of clinical psychiatry. 1998;59(20):22–33.

56. Oldfield RC. The assessment and analysis of handedness: the Edinburgh inventory. Neuropsychologia. 1971;9(1):97–113.

57. Canli T, Qiu M, Omura K, Congdon E, Haas BW, Amin Z, et al. Neural correlates of epigenesis. Proceedings of the National Academy of Sciences. 2006;103(43):16033–16038.

58. Patton JH, Stanford MS, Barratt ES. Factor structure of the Barratt impulsiveness scale. Journal of clinical psychology. 1995;51(6):768–774.

59. Horn W. Leistungsprüfsystem, LPS: Handanweisung für die Durchführung, Auswertung und Interpretation. Verlag Hogrefe; 1962.

60. Eriksen BA, Eriksen CW. Effects of noise letters upon the identification of a target letter in a nonsearch task. Perception & psychophysics. 1974;16(1):143–149.

61. Lang PJ, Bradley MM, Cuthbert BN, et al. International affective picture system (IAPS): Technical manual and affective ratings. NIMH Center for the Study of Emotion and Attention. 1997;1(39-58):3.

62. Luce RD, et al. Response times: Their role in inferring elementary mental organization. 8. Oxford University Press on Demand; 1986.

63. Hervey AS, Epstein JN, Curry JF, Tonev S, Eugene Arnold L, Keith Conners C, et al. Reaction time distribution analysis of neuropsychological performance in an ADHD sample. Child Neuropsychology. 2006;12(2):125–140.

64. Papaspiliopoulos O, Roberts GO, Sköld M. A general framework for the parametrization of hierarchical models. Statistical Science. 2007; p. 59–73.

65. Betancourt M, Girolami M. Hamiltonian Monte Carlo for hierarchical models. Current trends in Bayesian methodology with applications. 2015;79(30):2–4.

66. Bates D, Mächler M, Bolker B, Walker S. Fitting Linear Mixed-Effects Models Using lme4. Journal of Statistical Software. 2015;67(1):1–48. doi:10.18637/jss.v067.i01.

67. Windhoff M, Opitz A, Thielscher A. Electric field calculations in brain stimulation based on finite elements: an optimized processing pipeline for the generation and usage of accurate individual head models. Wiley Online Library; 2013.

68. Saturnino GB, Madsen KH, Thielscher A. Electric field simulations for transcranial brain stimulation using FEM: an efficient implementation and error analysis. Journal of neural engineering. 2019;16(6):066032.

69. Dale AM, Fischl B, Sereno MI. Cortical surface-based analysis: I. Segmentation and surface reconstruction. Neuroimage. 1999;9(2):179–194.

70. Fischl B, Sereno MI, Dale AM. Cortical surface-based analysis: II: inflation, flattening, and a surface-based coordinate system. Neuroimage. 1999;9(2):195–207.

71. Smith SM. Fast robust automated brain extraction. Human brain mapping. 2002;17(3):143–155.

72. Gabriel S, Lau R, Gabriel C. The dielectric properties of biological tissues: II. Measurements in the frequency range 10 Hz to 20 GHz. Physics in medicine & biology. 1996;41(11):2251.

73. Birot G, Spinelli L, Vulliémoz S, Mégevand P, Brunet D, Seeck M, et al. Head model and electrical source imaging: a study of 38 epileptic patients. NeuroImage: Clinical. 2014;5:77–83.

74. Oostenveld R, Fries P, Maris E, Schoffelen JM. FieldTrip: open source software for advanced analysis of MEG, EEG, and invasive electrophysiological data. Computational intelligence and neuroscience. 2011;2011.

75. Sejnowski TJ. Independent component analysis of electroencephalographic data. In: Advances in Neural Information Processing Systems 8: Proceedings of the 1995 Conference. vol. 8. MIT press; 1996. p. 145.

76. Chaumon M, Bishop DV, Busch NA. A practical guide to the selection of independent components of the electroencephalogram for artifact correction. Journal of neuroscience methods. 2015;250:47–63.

77. Ehinger BV, König P, Ossandón JP. Predictions of visual content across eye movements and their modulation by inferred information. Journal of Neuroscience. 2015;35(19):7403–7413.

78. Kriegeskorte N, Simmons WK, Bellgowan PS, Baker CI. Circular analysis in systems neuroscience: the dangers of double dipping. Nature neuroscience. 2009;12(5):535–540.

79. Maris E, Oostenveld R. Nonparametric statistical testing of EEG-and MEG-data. Journal of neuroscience methods. 2007;164(1):177–190.

80. Jeffreys D, Axford J. Source locations of pattern-specific components of human visual evoked potentials. I. Component of striate cortical origin. Experimental brain research. 1972;16(1):1–21.

81. Di Russo F, Martínez A, Sereno MI, Pitzalis S, Hillyard SA. Cortical sources of the early components of the visual evoked potential. Human brain mapping. 2002;15(2):95–111.

82. Gross J, Kujala J, Hämäläinen M, Timmermann L, Schnitzler A, Salmelin R. Dynamic imaging of coherent sources: studying neural interactions in the human brain. Proceedings of the National Academy of Sciences. 2001;98(2):694–699.

83. Percival DB, Walden AT, et al. Spectral analysis for physical applications. cambridge university press; 1993.

84. Brookes MJ, Vrba J, Robinson SE, Stevenson CM, Peters AM, Barnes GR, et al. Optimising experimental design for MEG beamformer imaging. Neuroimage. 2008;39(4):1788–1802.

85. Huang MX, Shih J, Lee R, Harrington D, Thoma R, Weisend M, et al. Commonalities and differences among vectorized beamformers in electromagnetic source imaging. Brain topography. 2004;16(3):139–158.

86. Helbling S. Advances in MEG methods and their applications to investigate auditory perception [doctoralthesis]. Universitätsbibliothek Johann Christian Senckenberg; 2017.

87. Gross J, Baillet S, Barnes GR, Henson RN, Hillebrand A, Jensen O, et al. Good practice for conducting and reporting MEG research. Neuroimage. 2013;65:349–363.

88. King JR, Dehaene S. Characterizing the dynamics of mental representations: the temporal generalization method. Trends in cognitive sciences. 2014;18(4):203–210.

89. Hebart MN, Baker CI. Deconstructing multivariate decoding for the study of brain function. Neuroimage. 2018;180:4–18.

90. Guggenmos M, Sterzer P, Cichy RM. Multivariate pattern analysis for MEG: A comparison of dissimilarity measures. NeuroImage. 2018;173:434–447.

91. Chang CC, Lin CJ. LIBSVM: a library for support vector machines. ACM transactions on intelligent systems and technology (TIST). 2011;2(3):1–27.

92. Dhamala M, Rangarajan G, Ding M. Analyzing information flow in brain networks with nonparametric Granger causality. Neuroimage. 2008;41(2):354–362.

93. Wang X, Chen Y, Bressler SL, Ding M. Granger causality between multiple interdependent neurobiological time series: blockwise versus pairwise methods. International journal of neural systems. 2007;17(02):71–78.

94. Salvatier J, Wiecki TV, Fonnesbeck C. Probabilistic programming in Python using PyMC3. PeerJ Computer Science. 2016;2:e55. doi:10.7717/peerj-cs.55.

95. Kruschke JK. Bayesian analysis reporting guidelines. Nature Human Behaviour. 2021;5(10):1282–1291.

96. Van Rossum G, Drake FL. Python 3 Reference Manual. Scotts Valley, CA: CreateSpace; 2009.

97. Anaconda Software Distribution; 2020. Available from: https://docs.anaconda.com/.

98. Barr DJ. Random effects structure for testing interactions in linear mixed-effects models. Frontiers in psychology. 2013;4:328.

99. R Core Team. R: A Language and Environment for Statistical Computing; 2018. Available from: https://www.R-project.org/.

100. Bates D, Sarkar D, Bates MD, Matrix L. The lme4 package. R package version. 2007;2(1):74.

